# Confounder-aware foundation modeling for accurate phenotype profiling in cell imaging

**DOI:** 10.1101/2024.12.23.630105

**Authors:** Giorgos Papanastasiou, Pedro P. Sanchez, Argyrios Christodoulidis, Guang Yang, Walter Hugo Lopez Pinaya

## Abstract

Image-based profiling is rapidly transforming drug discovery, offering unprecedented insights into cellular responses. However, experimental variability hinders accurate identification of mechanisms of action (MoA) and compound targets. Existing methods commonly fail to generalize to novel compounds, limiting their utility in exploring uncharted chemical space. To address this, we present a confounder-aware foundation model integrating a causal mechanism within a latent diffusion model, enabling the generation of balanced synthetic datasets for robust biological effect estimation. Trained on over 13 million Cell Painting images and 107 thousand compounds, our model learns robust cellular phenotype representations, mitigating confounder impact. We achieve state-of-the-art MoA and target prediction for both seen (0.66 and 0.65 ROC-AUC) and unseen compounds (0.65 and 0.73 ROC-AUC), significantly surpassing real and batch-corrected data. This innovative framework advances drug discovery by delivering robust biological effect estimations for novel compounds, potentially accelerating hit expansion. Our model establishes a scalable and adaptable foundation for cell imaging, holding the potential to become a cornerstone in data-driven drug discovery.

## Introduction

Image-based profiling is revolutionizing early-stage drug discovery, offering unprecedented capabilities to unravel the intricate cellular responses elicited by experimental (compound or genetic) perturbations [1]. This process involves treating cells with experimental perturbations and capturing the resulting morphological variations through cell microscopy [2]. Cell Painting (CP), the most widely established image-based profiling technique, utilizes six different dyes to image key cellular organelles and components, including RNA, DNA, mitochondria, plasma membrane, endoplasmic reticulum, actin cytoskeleton, and the Golgi apparatus [3, 4]. However, the specialized equipment and expertise required for CP are not readily available in many wet labs, highlighting the need for a more accessible foundation model for image-based profiling. Such a model could substantially accelerate drug discovery [1, 5, 6], by providing a valuable tool to explore the effects of novel chemical compounds on cell morphology, even in laboratories without specialized CP capabilities.

A central goal of CP is to quantitatively characterize compound mechanisms of action (MoA), providing fundamental insights into its biological activity and guiding the development of innovative therapeutics [5]. MoA identification remains challenging, often requiring a multi-faceted approach integrating data from various experimental technologies, such as transcriptomics, bioactivity assessments and CP [5]. Moreover, CP has shown promise in characterizing MoA [5], and developing computational methods to improve this process can enhance the efficiency and cost-effectiveness of early drug discovery [5, 6].

To characterize MoAs from CP images, automated image analysis focuses on quantifying the cellular morphology variations in response to experimental perturbations [1, 7]. Conventional image analysis methods rely on hand-engineered features capturing different aspects of cellular size, shape, intensity and texture across the CP stains [1, 5, 7]. Deep learning methods have outperformed non-batch-corrected conventional methods in MoA prediction [5], within focused [8–12] and broader MoA sets [13]. Other deep learning studies have recently demonstrated that integrating CP data with gene expression and chemical structure information enhances compound activity prediction [14], while weakly supervised learning can improve treatment identification [15]. In a recent study, a conditional generative adversarial network (GAN) was developed to reproduce cell phenotype changes induced by a small set of compound treatments [16]. The authors hypothesized that image synthesis would reduce cell-to-cell natural variability, which in turn would enhance the detection of subtle, treatment-induced morphological changes. Other efforts have focused on evaluating the capacity of generative models to synthesize CP-stained images from brightfield microscopy images (lacking CP dyes) [17–20]. Nevertheless, CP data and existing methods for estimating biological effects, such as MoA or target identification, are susceptible to substantial bias from confounders. These confounders encompass extraneous factors—including variations in laboratory conditions, experimental procedures, and imaging techniques—that can influence both the observed cellular morphologies and the measured biological effects, leading to spurious associations and obscuring true compound-induced phenotypic responses [5, 6]. These biases, arising from sources such as lab equipment variations, batch inconsistencies, well position effects, and other uncontrollable experimental factors, can significantly confound downstream analyses of CP data [5, 6]. Existing batch effect correction methods, such as Harmony, effectively account for batch effects and show promise for improving biological effect estimation from CP data [21]. However, these approaches typically assume that sources of variation can be captured through linear transformations. Despite the demonstrated capacity of generative models to synthesize CP data [16–20], no prior work has explicitly corrected for confounding biases in the CP image domain, potentially hindering the development of generative models for broader biological effect applications, such as predicting MoA and compound targets.

In our work, we developed a novel latent diffusion model (LDM)-based foundation model, pre-trained on a vast and diverse dataset comprising a total of 13,361,250 CP images corresponding to 107,289 compounds, from the Joint Undertaking for Morphological Profiling-Cell Painting Consortium (JUMP-CP) [6]. This model aims to learn generalizable representations of cellular phenotypic responses to compound perturbations, enabling it to potentially perform a wide range of downstream tasks in drug discovery, with minimal or no task-specific fine-tuning. To achieve this, we incorporated known confounding variables, i.e., source (laboratory), batch and well position, into the LDM architecture, effectively embedding a structural causal model (SCM) within the image generation process [22–25]. By explicitly encoding causal relationships, SCM-informed generative modeling can control for confounders and account for the complex interplay of causal factors in the image generation process [22, 26, 27]. Moreover, to inform the model about compound-specific effects, we incorporated chemical compound structure embeddings, derived by encoding Simplified Molecular-Input Line-Entry System (SMILES) representations using a pretrained MolT5 framework [28], as conditioning factors in the LDM. Therefore, our approach exposed the model to a vast and diverse spectrum of compound-induced morphological changes while simultaneously accounting for their underlying causal factors (confounders).

We investigate our confounder-aware (SCM-conditioned) foundation model’s ability for MoA identification and compare its performance against real, real batch-corrected JUMP-CP data and a similar foundation model lacking the incorporated SCM. We conduct the same evaluation for compound target prediction, a task not previously investigated by generative modeling or classification techniques in the CP domain. This is particularly important for broadly evaluating the biological effects of chemical compounds, as identifying potential targets provides crucial insights into the specific mechanisms by which these compounds exert their effects, complementing MoA prediction. We demonstrate the capacity of our confounder-aware foundation model for compound-to-image synthesis and its ability to improve both MoA and compound target identification. This evaluation encompasses two distinct scenarios: assessing the model’s performance in MoA and target identification using synthetic images generated from a) compounds included in the training dataset and b) novel compounds unseen during training. This latter scenario provides compelling evidence for the model’s ability to extrapolate beyond the training data, demonstrating its generalization capabilities and unlocking the potential for robust predictions within uncharted chemical spaces. Therefore, we present a novel confounder-aware foundation model that improves the accuracy of MoA and target prediction from CP images, demonstrating strong generalization to novel compounds and paving the way for enhanced hit expansion and a deeper understanding of compound-induced cellular responses.

## Results

This study involved developing a confounder-aware LDM foundation model and evaluating its ability to predict MoA and targets for both seen and unseen compounds. Furthermore, we qualitatively and quantitatively assessed the model’s impact on separating compounds, batch effects, and MoAs, while reducing within-image variability. We extensively compared the confounder-aware model against real images, batch-corrected images, and a non-confounder-aware model.

To explore the impact of scaling confounding factor combinations (N=10 versus 100) on MoA and target prediction for seen and unseen compounds, we synthesized two sets of CP images using a g-estimation-inspired methodology [29]. The synthesized images consisted of a smaller set with 5,000 images (N=10) and a larger set with 50,000 images (N=100). These images were then used to generate cell profiles, which were subsequently evaluated for their ability to predict MoA and compound targets. For the non-confounder-aware model, we generated the same number of images to ensure fair comparisons during evaluation.

### Synthetic image generation with a confounder-aware LDM

Our confounder-aware LD foundation model, incorporating MolT5-derived chemical embeddings and trained to disentangle and control for confounding factors, generated synthetic CP images that accurately captured compound-induced morphological changes. The process of generating synthetic CP images using our confounder-aware LDM, incorporating MolT5 embeddings and controlling for confounding factors, is depicted in Figure 1.

**Figure 1).**
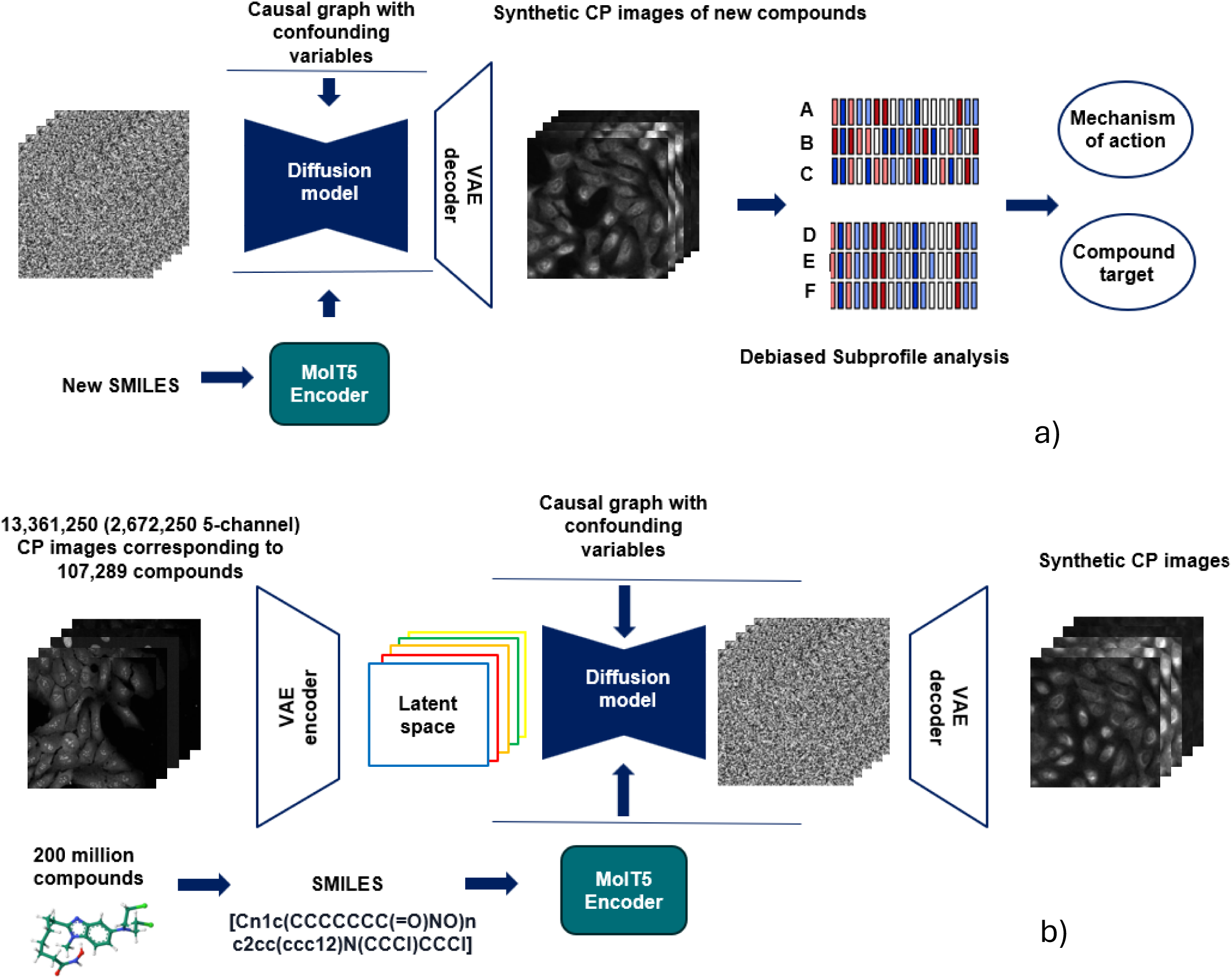
(a) Our novel confounder-aware foundation model framework. (a) A SCM-conditioned latent diffusion model was trained using a total of 13,361,250 CP images (2,672,250 5-channel CP images). Each 5-channel set had multiple field of views, and were typically acquired across ≥ 1 sources, and multiple batches, plates and well positions. The SCM-conditioned latent diffusion model learns the effects of the biases on the CP images, for each of the 107,289 chemical compounds used as inputs. The SMILE notation of each chemical compound was ran through a MolT5 Transformer that it was pretrained on 200 million compounds [28]. (b) Novel molecules, represented in SMILE notation, are encoded and input to the trained foundation model. This model, conditioned on confounders (source, batch, well position) and sampled Gaussian noise, aims to generate synthetic CP images that account for these confounders, ultimately allowing to debias downstream tasks (subprofile analysis for MoA and target identification) through a g-estimation-based method (see Methods) [29]. SCM: structural causal model, CP: cell painting, SMILE: Simplified Molecular Input Line Entry System, MoA: mechanism of action.

### Exploratory analysis and causal modeling of confounders

To explore the complex relationships and potential confounding factors within the JUMP-CP dataset, we conducted exploratory analyses of the distributions of compounds across sources, batches, plates, and wells. The Sankey diagram in Figure 2a visualizes these relationships of 5 randomly selected compounds, highlighting the interconnectedness of these factors.

To address potential confounding biases, we developed a causal graph (Figure 2b) and employed a g-estimation-based method to control for confounders in the synthetic image generation process (Figure 2c), ensuring that the generated images accurately reflect the causal relationships between compounds, phenotypes, and confounding factors.

**Figure 2).**
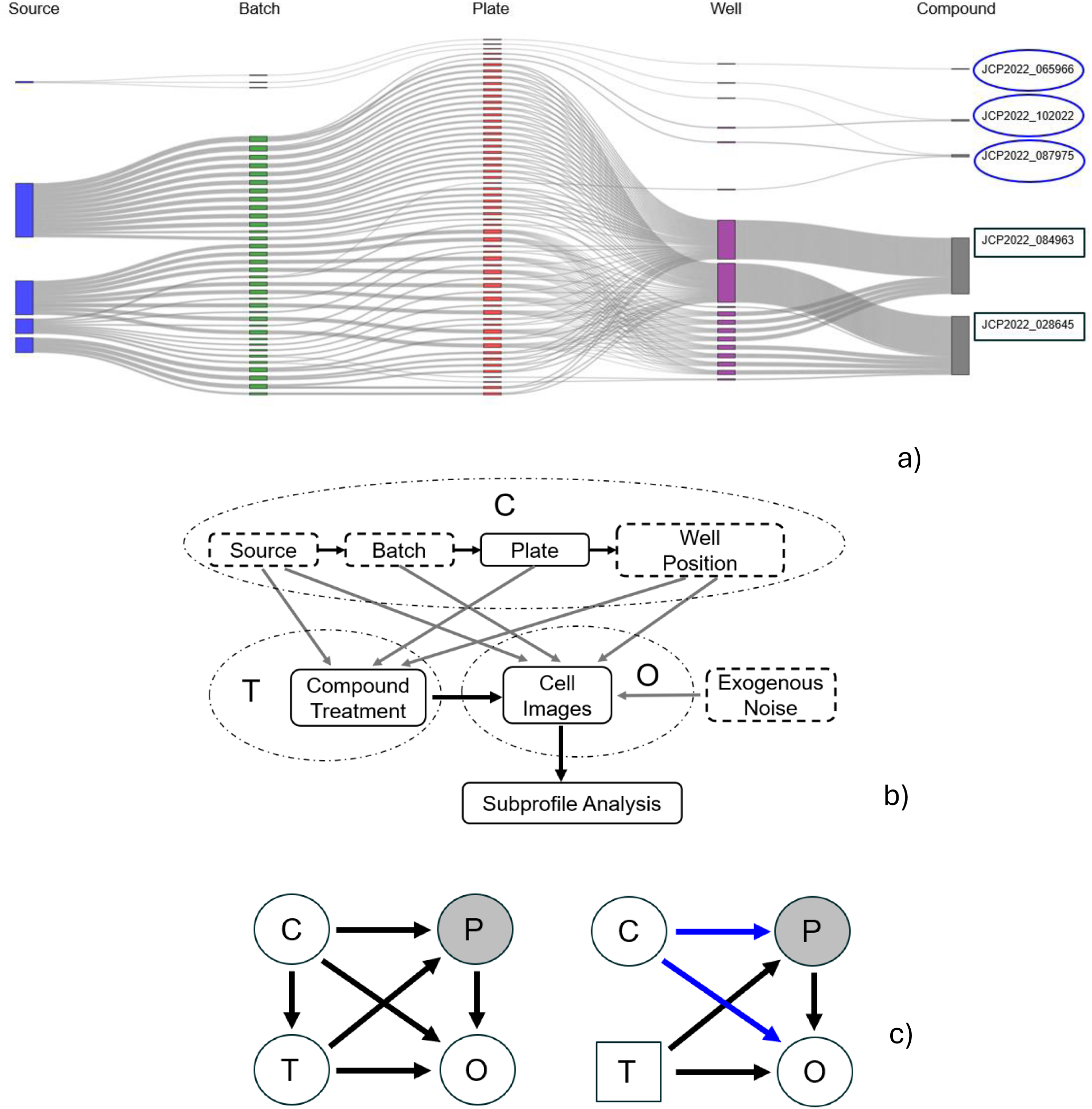
(a) Exploratory analysis: Sankey diagram showing the distributions of 5 randomly selected compounds across sources, batch, plates, and wells in the JUMP-CP dataset. Blue ovals indicate compounds with infrequent distribution, while black rectangles represent those with frequent occurrence across sources, batches, plates, and well positions. (b) Detailed causal graph showing the causal paths between each of the known confounders C, compound treatment T and images O. To improve clarity, unobserved phenotypes P are not shown here. (c) Left: causal graph showing the causal paths between the confounders C, compound treatment T, true phenotype P (unobserved in the data), and image observations O, in the real distribution. Right: Same causal graph in the synthetic distribution, after adjusting for confounders using a g-estimation-based method (shown with blue, see details in Methods).

### Qualitative and quantitative evaluation of confounder-aware image generation

Initially, to assess the impact of our confounder-aware foundation model, we performed a qualitative evaluation of its ability to isolate the effect of confounders. As shown in Figure 3, we generated synthetic samples of the same compound (DMSO and AMG-900) under different batch and well position conditions, using a single random seed to minimize variability in cell position and density across confounder sets [16]. These synthetic samples were then compared against real data with matching conditions. Our model effectively minimized within-image variability while capturing across-batch and –well position variability, demonstrating its ability to synthesize images conditioned on the observable batch and well position effects.

**Figure 3).**
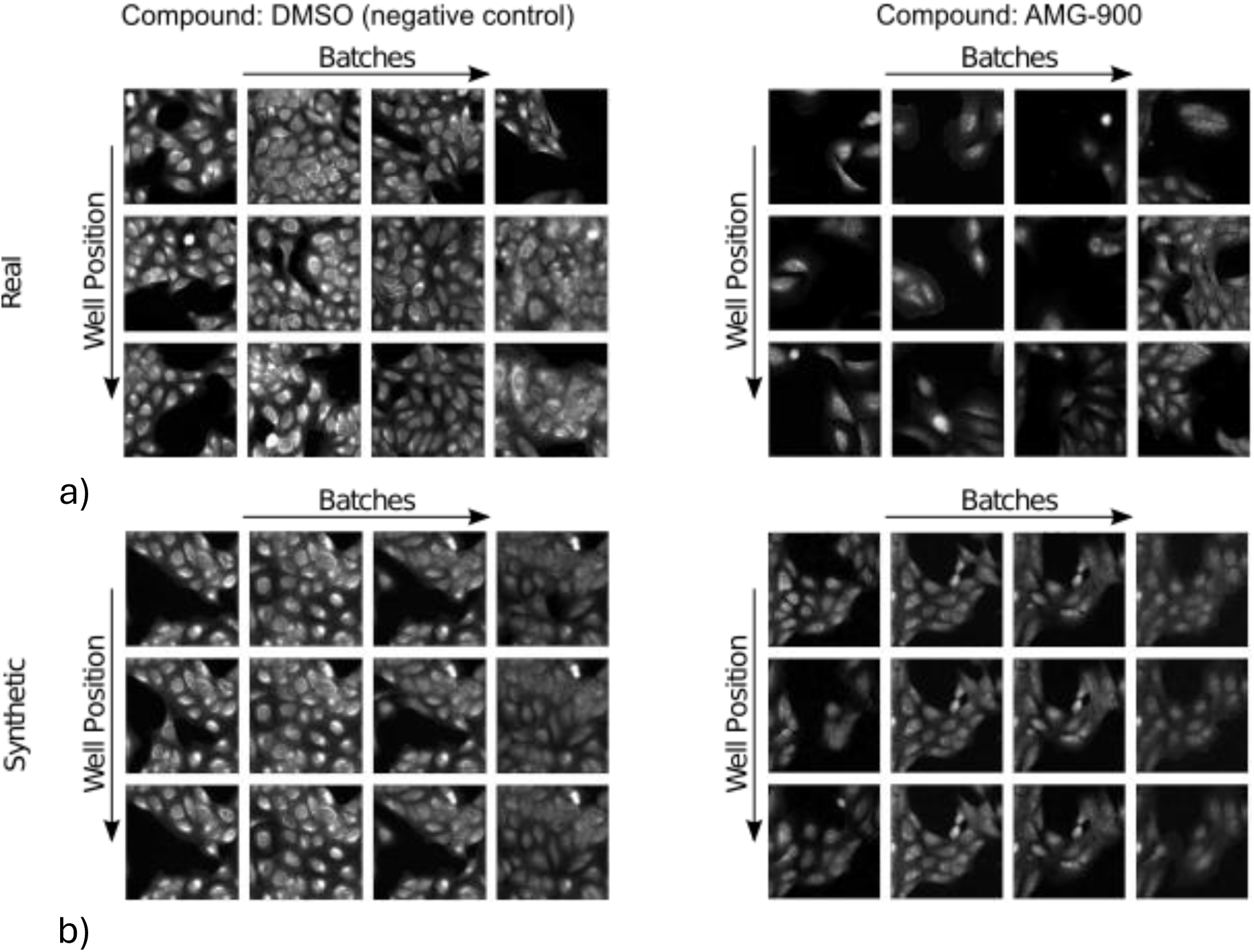
Comparison of real (a) and synthetic (b) CP images across varying experimental conditions. The top row displays real images, while the bottom row shows synthetic images generated by the confounder-aware foundation model. Images are presented for both DMSO (negative control) and the AMG-900 compound (MoA: aurora kinase inhibitor), across different batches and well positions. Notably, it is observed that the confounder-aware foundation model can minimize within-image variability, while increasing (capturing) across-batch variability since it synthesizes images conditioned on the observable batch effect. Synthetic images were generated using a single random seed to reduce undesired variability in cell position and density. CP: cell painting, MoA: mechanism of action.

Subsequently, to gain an initial quantitative understanding of how confounders influence cell profiles and compound-induced morphological changes, we performed a UMAP analysis, visualizing the clustering patterns of CP image-derived cell profiles in Figure 4. Each blue circle in panels a-f, represents cell profiles from a 5-channel CP image set with a unique combination of confounders (source, batch, well position), while panels g-i show cell profiles aggregated across all confounder combinations for each compound. Clear distinctions are observed between the clusters derived from real data, confounder-aware foundation model data, and non-confounder-aware foundation model data, highlighting the differing influence of confounders across the datasets.

**Figure 4).**
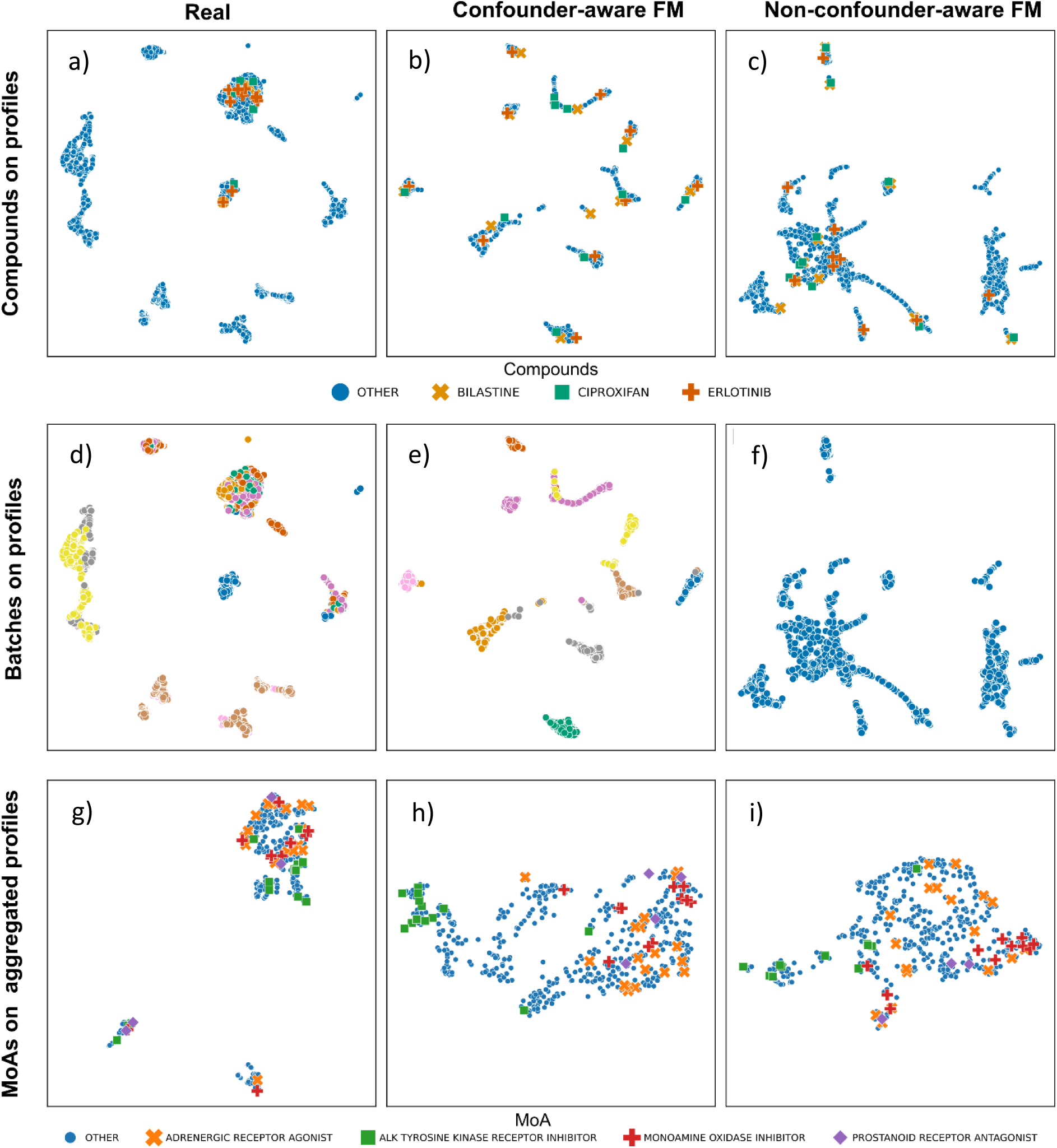
UMAP analysis reveals the impact of confounder-aware foundation model (FM) image generation on cluster separation. Clusters across panels reflect the influence of confounders on cell profiles. Each blue circle in panels a-f represents cell profiles from a 5-channel CP image set with a unique combination of confounders (source, batch, well position). Panels g-i show cell profiles aggregated across all confounder combinations for each compound. Confounder-aware FM (panels b, e, h; generated using a g-estimation-based approach [29], see Methods) improves separation of compounds (a-c), batch effects (d-f), and MoAs (g-i) compared to Real and non-confounder-aware FM, highlighting its ability to mitigate confounder influence. UMAP: uniform manifold approximation and projection, MoA: mechanism of action. In panels d-f, each color corresponds to a different batch. Non-confounder-aware FM is confounder-agnostic; thus, no batch effects are shown in panel f.

To assess compound separation, we visualized 3 randomly selected compounds (Bilastine, Ciproxifan, Erlotinib) across all datasets (Figure 4a-c). The confounder-aware foundation model (Figure 4b) achieved clear separation of these compounds, while the real data and non-confounder-aware model (Figure 4a and c) showed substantial overlap within the observed clusters. Similarly, visualization of batch effects (Figure 4d-f) revealed that the confounder-aware model (Figure 4e) effectively separated batches, whereas the real data and non-confounder-aware model (Figure 4d and f) again exhibited overlap. Furthermore, evaluation of MoAs on aggregated cell profiles (Figure 4g-i) showed that the confounder-aware foundation model (Figure 4h) demonstrates a clearer separation of MoAs compared to the real data and the non-confounder-aware model (Figure 4g and i). This suggests that the model effectively accounts for and mitigates the influence of confounders, allowing for a more accurate representation of MoA-related differences.

### Confounder-aware foundation model improves MoA prediction accuracy

Firstly, to evaluate the impact of our confounder-aware foundation model on MoA identification for compounds seen during training, but with unseen images (i.e., images corresponding to batches not encountered during training), we compared its performance to three other settings: (1) the original real CP image data, (2) real batch-corrected CP image data (using Harmony), and (3) synthetic data generated by a non-confounder-aware model. In each setting, we used a nearest neighbor classifier to assess the accuracy of the estimated Biosimilarities in identifying known MoAs from the Drug Repurposing Hub.

Figure 5a shows improved mAP [30] and ROC-AUC [31] performance for the confounder-aware model in identifying MoAs. With 50,000 images (N=100 confounder combinations) and 5,000 images (N=10) for the confounder-aware model, the mAP was 0.08 and 0.07, respectively. The non-confounder-aware model had a mAP of 0.07 for both 50,000 and 5,000 images. Real and real batch-corrected data had mAPs of 0.06 and 0.07, respectively.

**Figure 5).**
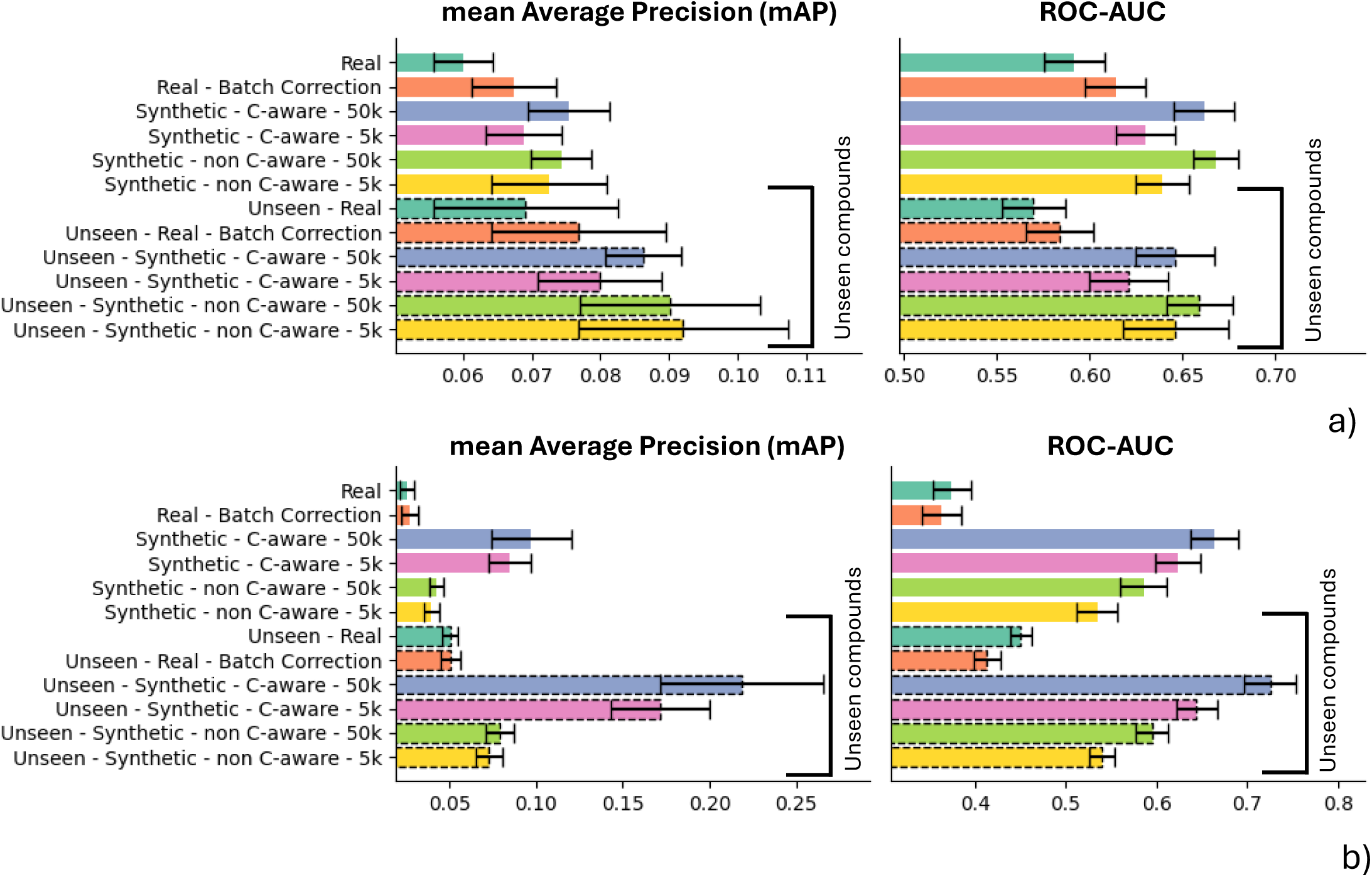
Performance comparisons for MoA (a) and compound target (b) identification across real, real batch-corrected, confounder-aware (SCM-conditioned) foundation model-derived synthetic, and non-confounder-aware (SCM-free) foundation model-derived synthetic data. Error bars represent the standard deviation across 10-fold cross-validation, obtained by varying the selection of the compounds used to derive the “reference subprofile” across folds.

With 50,000 images (N=100) and 5,000 images (N=10) for the confounder-aware model, the ROC-AUC was 0.66 and 0.63, respectively. The non-confounder-aware model had a ROC-AUC of 0.64 for both 50,000 and 5,000 images. Real and real batch-corrected data had ROC-AUCs of 0.59 and 0.61, respectively.

ANOVA analysis with Tukey’s honestly significant difference (HSD) post-hoc test [32] comparing mAP results across all real and synthetic data types, revealed that both the confounder-aware and non-confounder-aware foundation models significantly outperformed real data in MoA prediction (Figure 5a, Table 1, Supplementary Data 1), with no other significant differences observed. Repeating the ANOVA with Tukey’s HSD test to compare ROC-AUC results across all data types, both foundation models significantly outperformed real and real-batch corrected data (Figure 5a, Table 1, Supplementary Data 1). No significant differences were observed between the two foundation models (comparing the same number of images: 50,000 versus 50,000 and 5,000 versus 5,000; Table 1).

**Table 1).**
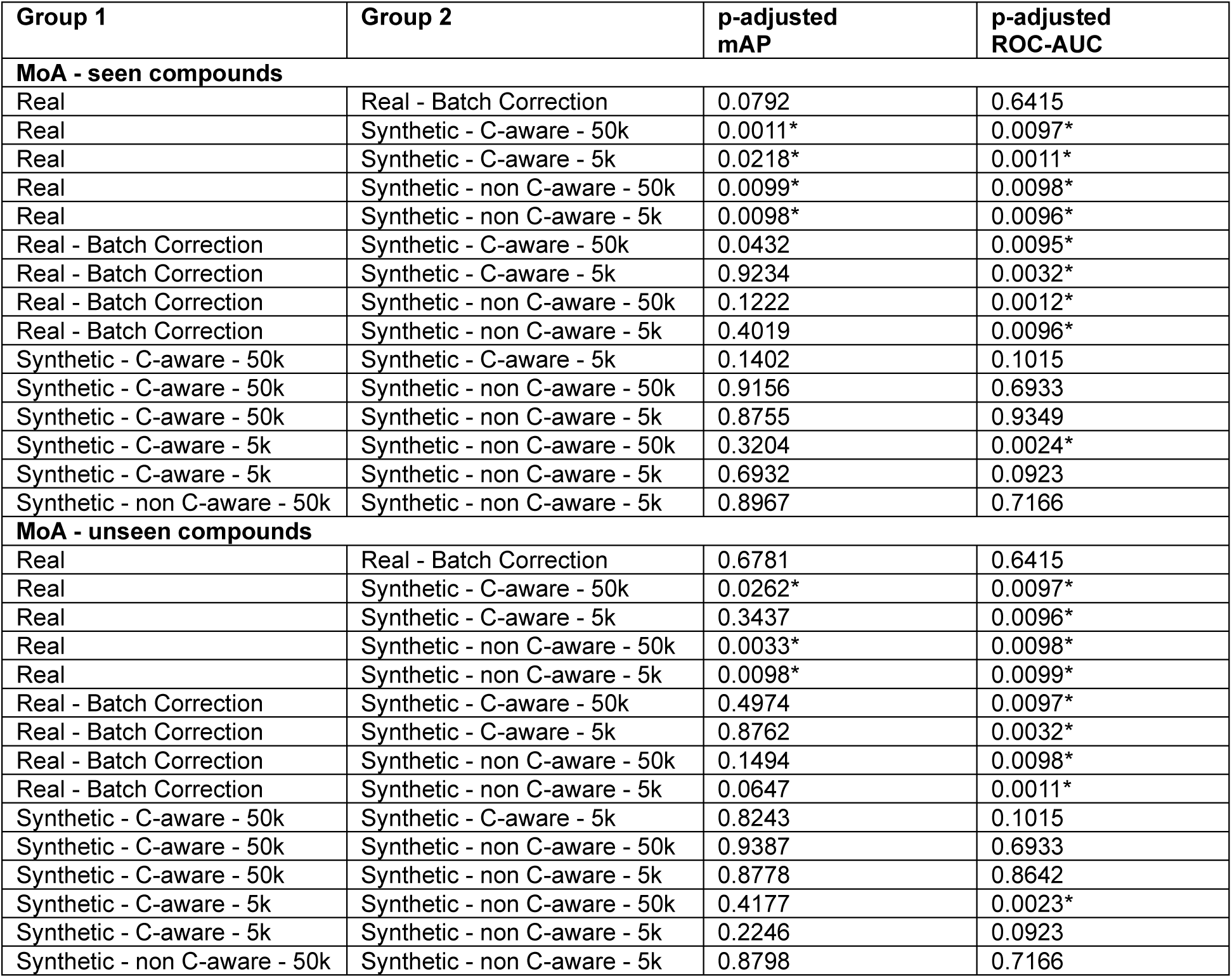
Assessing differences in MoA prediction performance across image types. Results of ANOVA with Tukey’s HSD post-hoc test comparing mean average precision (mAP) and area under the receiver operating characteristic curve (ROC-AUC) for MoA prediction across different image types: real, real batch-corrected, confounder-aware (C-aware) synthetic, and non-confounder-aware (non-C-aware) synthetic images. Comparisons were performed for both seen and unseen compounds during training. Adjusted p-values are presented. Statistical significance is indicated with *. MoA: mechanism of action; HSD: honestly significant difference; C-aware: confounder-aware.

### Confounder-aware foundation model improves target prediction accuracy

We conducted the same analysis for target prediction, using compounds present during training, but with unseen images. Figure 5b shows that the confounder-aware model consistently outperformed all other data types in predicting targets, as evidenced by the mAP and ROC-AUC results. With 50,000 images (N=100) and 5,000 images (N=10) for the confounder-aware model, the mAP was 0.10 and 0.08, respectively. The non-confounder-aware model had a mAP of 0.04 for both 50,000 and 5,000 images. Real and real batch-corrected data had mAPs of 0.02 and 0.03, respectively.

The confounder-aware model had a ROC-AUC of 0.65 and 0.62 with 50,000 (N=100) and 5,000 (N=10) images, respectively. The non-confounder-aware model had a ROC-AUC of 0.66 and 0.65 for those respective image sets. Real and real batch-corrected data had ROC-AUCs of 0.57 and 0.58, respectively.

ANOVA with Tukey’s HSD post-hoc test revealed that the confounder-aware model significantly outperformed all other image types in both mAP and ROC-AUC (Figure 5b, Table 2, Supplementary Data 1). The non-confounder-aware model has also significantly outperformed the real and real batch-corrected data, but underperformed against the confounder-aware model.

**Table 2).** Assessing differences in compound target prediction performance across image types. Results of ANOVA with Tukey’s HSD post-hoc test comparing mean average precision (mAP) and area under the receiver operating characteristic curve (ROC-AUC) for target prediction across different image types: real, real batch-corrected, confounder-aware (C-aware) synthetic, and non-confounder-aware (non-C-aware) synthetic images. Comparisons were performed for both seen and unseen compounds during training. Adjusted p-values are presented. Statistical significance is indicated with *. HSD: honestly significant difference; C-aware: confounder-aware.

| Group 1 | Group 2 | p-adjusted mAP | p-adjusted ROC-AUC |
| --- | --- | --- | --- |
| **Target - seen compounds** |  |  |  |
| Real | Real - Batch Correction | 0.8968 | 0.8806 |
| Real | Synthetic - C-aware - 50k | 0.0011* | 0.0098* |
| Real | Synthetic - C-aware - 5k | 0.0098* | 0.0099* |
| Real | Synthetic - non C-aware - 50k | 0.0161* | 0.0096* |
| Real | Synthetic - non C-aware - 5k | 0.0823 | 0.0096* |
| Real - Batch Correction | Synthetic - C-aware - 50k | 0.0097* | 0.0097* |
| Real - Batch Correction | Synthetic - C-aware - 5k | 0.0096* | 0.0098* |
| Real - Batch Correction | Synthetic - non C-aware - 50k | 0.0424* | 0.0099* |
| Real - Batch Correction | Synthetic - non C-aware - 5k | 0.1789 | 0.0096* |
| Synthetic - C-aware - 50k | Synthetic - C-aware - 5k | 0.1705 | 0.0098* |
| Synthetic - C-aware - 50k | Synthetic - non C-aware - 50k | 0.0099* | 0.0097* |
| Synthetic - C-aware - 50k | Synthetic - non C-aware - 5k | 0.0011* | 0.0099* |
| Synthetic - C-aware - 5k | Synthetic - non C-aware - 50k | 0.0010* | 0.0098* |
| Synthetic - C-aware - 5k | Synthetic - non C-aware - 5k | 0.0098* | 0.0099* |
| Synthetic - non C-aware - 50k | Synthetic - non C-aware - 5k | 0.9312 | 0.0096* |
| **Target - unseen compounds** |  |  |  |
| Real | Real - Batch Correction | 0.9431 | 0.0098* |
| Real | Synthetic - C-aware - 50k | 0.0011* | 0.0099* |
| Real | Synthetic - C-aware - 5k | 0.0010* | 0.0011* |
| Real | Synthetic - non C-aware - 50k | 0.0831 | 0.0012* |
| Real | Synthetic - non C-aware - 5k | 0.2753 | 0.0010* |
| Real - Batch Correction | Synthetic - C-aware - 50k | 0.0099* | 0.0010* |
| Real - Batch Correction | Synthetic - C-aware - 5k | 0.0098* | 0.0098* |
| Real - Batch Correction | Synthetic - non C-aware - 50k | 0.0914 | 0.0097* |
| Real - Batch Correction | Synthetic - non C-aware - 5k | 0.2962 | 0.0098* |
| Synthetic - C-aware - 50k | Synthetic - C-aware - 5k | 0.0012* | 0.0099* |
| Synthetic - C-aware - 50k | Synthetic - non C-aware - 50k | 0.0011* | 0.0097* |
| Synthetic - C-aware - 50k | Synthetic - non C-aware - 5k | 0.0010* | 0.0098* |
| Synthetic - C-aware - 5k | Synthetic - non C-aware - 50k | 0.0098* | 0.0099* |
| Synthetic - C-aware - 5k | Synthetic - non C-aware - 5k | 0.0097* | 0.0098* |
| Synthetic - non C-aware - 50k | Synthetic - non C-aware - 5k | 0.8976 | 0.0096* |

### Generalizing MoA prediction to unseen compounds

To assess model generalizability, we conducted the same MoA prediction analysis using novel compounds unseen during training, observing similar trends as with seen compounds (Figure 5a).

The confounder-aware model had a mAP of 0.09 with 50,000 images (N=100) and 0.08 with 5,000 images (N=10). The non-confounder-aware model had a mAP of 0.09 for both image sets. Real and real batch-corrected data had mAPs of 0.07 and 0.08, respectively.

The confounder-aware model had an ROC-AUC of 0.65 and 0.62 with 50,000 (N=100) and 5,000 (N=10) images, respectively. The non-confounder-aware model had a ROC-AUC of 0.66 and 0.65 for those respective image sets. Real and real batch-corrected data had ROC-AUCs of 0.57 and 0.58, respectively.

ANOVA with Tukey’s HSD post-hoc test revealed that both foundation models significantly outperformed real data in MoA prediction based on both mAP and ROC-AUC (Figure 5a, Table 1, Supplementary Data 1). For ROC-AUC, both models also significantly outperformed real batch-corrected data. No significant differences were observed between the two foundation models (comparing the same number of images: 50,000 versus 50,000 and 5,000 versus 5,000; Table 1).

### Generalizing target prediction to unseen compounds

When predicting targets for unseen compounds, the confounder-aware model achieved a mAP of 0.22 with 50,000 images (N=100) and 0.17 with 5,000 images (N=10) (Figure 5b). The non-confounder-aware model had a mAP of 0.08 and 0.07, for those respective image sets. Real and real batch-corrected data had a mAP of 0.05.

The confounder-aware model had ROC-AUCs of 0.73 and 0.64 with 50,000 (N=100) and 5,000 (N=10) images, respectively. The non-confounder-aware model had ROC-AUCs of 0.59 and 0.54 with those respective image sets. Real and real batch-corrected data had ROC-AUCs of 0.45 and 0.41, respectively.

ANOVA with Tukey’s HSD post-hoc test revealed that the confounder-aware model significantly outperformed all other image types in both mAP and ROC-AUC (Figure 5b, Table 2, Supplementary Data 1). Supplementary Figures 1 and 2 show examples of MoAs and targets where the confounder-aware model outperformed the other approaches.

## Discussion

In this study, we introduced a confounder-aware latent diffusion-based foundation model to address the challenge of confounding factors in image-based profiling for drug discovery, as well as the unmet need of generalizing to unseen compounds. Our model, trained on a vast CP dataset, generated synthetic images that accurately captured compound-induced morphological changes while mitigating the impact of confounders. This resulted in significantly improved and debiased MoA and target prediction for both seen and unseen compounds, as demonstrated by increased mAP and ROC-AUC and confirmed by ANOVA with Tukey’s HSD post-hoc tests.

This study presents, to our knowledge, the first large-scale generative model for cell imaging rigorously evaluated for its ability to predict MoAs and compound targets. Τhis evaluation includes the prediction of compound targets, a task not previously addressed by generative modeling [16–20] or classification techniques [8–13], applied to CP data. This is particularly valuable for broadly evaluating the biological effects of chemical compounds, as target identification provides crucial mechanistic insights-ultimately revealing what the compound aims to affect-complementing MoA prediction [1–7]. A rigorous evaluation of downstream tasks such as MoA and target prediction, and generalization to novel compounds, is essential to thoroughly assess the capacity of our foundation model to learn the underlying data generation process. Accurately predicting MoA and targets is inherently challenging due to the complex interplay between compound actions (e.g., inhibition, antagonist, agonist, or kinase action) and their effects on targets (e.g., proteins, enzymes, molecules) [1, 5]. It is also known that MoA annotation is inherently limited by factors such as dose-dependency, variability in dose-response curves, and the potential for polypharmacology [5]. A previous work has explored the use of contrastive learning to enable cross-modal queries between chemical structures and CP images [34]. Their method, termed CLOOME, utilized a dataset of approximately 760,000 CP images and 30,404 compounds, and demonstrated improved performance against non-batch-corrected data using ranking metrics. However, this approach did not explicitly address the learning of the underlying data generation process or incorporate mechanisms to account for confounding factors. As a retrieval-based technique, CLOOME’s performance may be influenced by the specific patterns in its training data and the extent of chemical and phenotypic diversity it encompasses. Using a relatively small CP dataset, another work used a conditional GAN to visualize and enhance understanding of how compounds influence cellular responses [16]. Our previous work developed and evaluated generative models for producing CP images from brightfield microscopy data [20, 35]. Our confounder-aware model demonstrates strong performance across seen and unseen compounds, and varying levels of confounding factors (N=10 to N=100) (Figures 5a and b, Tables 1 and 2). Moreover, our confounder-aware model showed statistically significant improvements in mAP compared to real data for both MoA and target prediction in seen and unseen compounds. This is the first study to benchmark these metrics at such a large scale, providing valuable insights into the performance of generative models for cell imaging. Therefore, unlike previous correlational techniques, our confounder-aware foundation model presents a unique and advanced approach, potentially enabling superior generalization and debiasing of downstream tasks.

Our confounder-aware model, by mapping over 107,000 compound-induced morphological changes in a causal manner, is capable of exploring both structure-based and biology-based hit expansion [33]. Structure-based hit expansion assumes smaller changes to the compound within a defined chemical space, while biology-based hit expansion is agnostic to chemical structure, focusing instead on biological effects and phenotypic profiles [33]. Accurate prediction of MoAs and targets for unseen compounds, achieved through our integrated Biosimilarity analysis, unlocks the potential to identify further novel MoAs and targets. Our model can be readily expanded with additional CP data, further refining its representation of the data generation process.

While our exploratory analysis (Figures 3-4) demonstrated improved separation of confounding effects, compounds, and MoAs, and our confounder-aware model showed significantly enhanced performance in target identification compared to all other data types, we observed comparable performance between the confounder-aware and non-confounder-aware models in MoA identification. This suggests that compound conditioning alone may be sufficient to guide image generation and support downstream analysis for MoA identification. Furthermore, the specific experimental confounders we considered may exert a greater influence on target identification, a novel downstream task in this context. It is important to acknowledge that our causal mechanism can be further refined by incorporating additional confounding factors known to influence MoA estimation, such as compound concentration and bioactivity, which were not available in the JUMP-CP dataset. Future work could explore the integration of external data and fine-tuning our model with these additional confounders to potentially further enhance performance.

The following limitations should be considered when interpreting our findings. First, we should acknowledge the influence of other potential confounders such as dose-dependency, variability in dose-response curves, and the potential for polypharmacology [1, 5]. Since our selected data lacked dose variation (10 μM across all sources), dose-response curves were unavailable, and all 525 compounds (used for MoA evaluation) had only 1 known MoA, we focused on accounting for the remaining experimental biases. Our confounder-aware model can be readily extended to include additional conditions in the SCM as they become available, enabling further refinements by our group and the broader research community. While deep learning techniques for MoA classification have shown promise [8–13], we used cell profiles to leverage Harmony for batch effect correction, which has demonstrated effectiveness in mitigating batch effects while preserving biological variability and interpretability [21]. We utilized 525 and 465 compounds for MoA and target prediction evaluation, respectively (Supplementary Table 2). This represents all annotated data from our JUMP-CP dataset found in the Drug Repurposing Hub, but it is a small subset of the overall JUMP-CP dataset. Future work could explore incorporating additional annotations or developing alternative evaluation strategies. Various batch correction techniques could be further explored to remove biases. Finally, alternative classification methods could be evaluated for biological effect estimation.

Our novel confounder-aware foundation model, with its demonstrated ability to accurately predict MoAs and targets even for novel compounds, establishes a new state-of-the-art and benchmark for cell imaging, with the potential to revolutionize hit expansion and deepen our understanding of compound-induced cellular responses.

## Methods

### Ethical statement

This study utilized exclusively computational methods with publicly available data, adhering to all relevant ethical principles for machine learning and data science research. No human or animal subjects, nor any wet-lab experimentation were performed. To broaden access, facilitate transparency and catalyze the use and deployment of our foundation model, we provide open access to our codebase.

### Data extraction and curation

Data extraction and curation details are summarized in the Supplementary Table 1. CP, the leading image-based profiling technique, utilizes six fluorescent dyes to label major cellular components: nucleus (DNA), endoplasmic reticulum, nucleoli, cytoplasmic RNA, actin, Golgi apparatus, plasma membrane, and mitochondria [1, 3, 5, 6, 14, 15, 21]. The JUMP-CP consortium, a collaborative effort led by the Broad Institute and involved various academic as well as industry partners and 12 pharmaceutical companies, has generated a large, optimized and diverse dataset to empower the development of advanced image analysis algorithms [6, 36]. JUMP-CP is a vast dataset comprising millions of CP images, profiling the morphological effects of >116,000 compounds across 4 distinct datasets: cpg0000, cpg0001, cpg0002 and cpg0016 [5, 6].

Following protocol optimization and standardization across three pilot datasets (cpg0000, cpg0001, cpg0002), the JUMP-CP consortium generated the largest principal dataset (cpg0016), comprising millions of 5-channel CP images corresponding to over 116,000 compounds from 12 sources (laboratories). Two foundation models were trained on the principal dataset: one incorporating confounder-awareness via SCM-conditioning (see “Defining the structural causal model”), and a second without this conditioning (SCM-free). Our dataset comprised all available data from 5 sources, consisting of 2,672,250 5-channel images (1 channel per fluorescent dye). The 5 sources (JUMP-CP laboratories 1-3, 9 and 11) were selected for their contribution of partially overlapping compound sets, ensuring that each compound was assayed using different instruments and microscopes across multiple laboratories (Figure 2a) [6]. To reduce cell variability, we focused on analyzing the U2OS cell line [6]. This yielded a total of 13,361,250 CP images, corresponding to 107,289 compounds which were organized into 48 batches and 832 plates. Each plate contains from 384 to 1536 wells, each imaged from 4 to 16 different fields of view with a microscope. Each well contained either a single chemical compound or a negative control (DMSO) in predefined positions [5, 6].

To evaluate our SCM-conditioned and SCM-free foundation models versus real data in the setting of MoA and target characterization, we retrieved all available MoA and target annotations for our selected compounds from the Broad Institute Drug Repurposing Hub http://www.broadinstitute.org/repurposing [37]. The Drug Repurposing Hub comprises a unique assemblage of compounds designed to facilitate drug repurposing efforts. This curated library encompasses a diverse range of >7,000 compounds with established clinical histories, including marketed drugs, those previously evaluated in human clinical trials, and preclinical tools [37]. All compounds were sourced from over 50 chemical vendors and subjected to rigorous purity validation, before inclusion in the library [37]. MoA characterization is a central goal in image-based cell profiling, offering a powerful framework for understanding compound effects [38]. However, it is known that source, batch, and well position confounders can bias or attenuate the compound effect, potentially affecting MoA and other biological effect characterizations [1, 5, 6]. Adjusting for the aforementioned confounders can potentially improve the accuracy of MoA characterization, which is important particularly for new or uncharacterized compounds [5]. To evaluate the accuracy of subprofile analysis in MoA and compound target identification from both the confounder-aware (SCM-conditioned) and non-confounder-aware (SCM-free) foundation model, against the real JUMP-CP, we used mean average precision and area under the receiver operating characteristic curve (see the subsection “Evaluation metrics”). This evaluation was conducted for both synthetic images generated from compounds included in the training dataset and those derived from compounds entirely novel to the model, enabling an assessment of the models’ generalization capabilities. From 107,289 compounds involved in our study (Figure 1), we identified 525 compounds with MoA and target annotations (used as ground truths) in the Drug Repurposing Hub library. These compounds were divided into 2 groups. The first group was used to establish “reference profiles” based on their MoA and target information (see subsection “MoA and target identification via subprofiling”). The second group, containing compounds both seen and unseen during model training, was then compared to these reference profiles, to derive Biosimilarity scores. This allowed us to evaluate the performance of our confounder-aware-model and non-confounder-aware-model-derived cell profiles in characterizing the effects of both familiar (seen during training) and novel (unseen) compounds. Further details are described in the “MoA and target identification via subprofiling” and “Evaluation metrics” subsections.

Each source (laboratory) of the JUMP-CP consortium conducts multiple batches of experiments, with each batch comprising multiple plates containing different compounds and negative controls in predefined well positions. A unique compound is applied to each well, but the selection of compounds varies across sources, batches and well positions. These variations (source, batch and well position) introduce potential biases, as illustrated in Figure 2a. The figure shows the uneven distribution of compounds across source, batch and well position combinations, highlighting the potential for systematic bias. Such technical variations can introduce systematic noise, potentially masking the true biological phenotypes [1, 3, 5, 6, 15, 21].

### Defining the structural causal model

We hypothesize that accurate estimation of compound effects from observational cell imaging data requires careful consideration of the causal relationships between known confounders, image generation, and the downstream task of interest i.e., MoA and target identification [23, 26, 27]. Figure 2b shows in detail the causal paths between the known confounders (source, batch, plate, and well position), the compound treatment, and the resulting cell images, all within the SCM structure. A SCM employs a directed acyclic graph (DAG) to represent causal relationships between the confounding variables. In the DAG structure, the nodes represent variables and edges indicate direct causal effects [24, 25]. SCM-informed generative modeling offer a powerful mechanism for capturing confounding factors within the image generation process [22, 26, 27]. The left panel of Figure 2c depicts the SCM that captures the causal relationships between confounders, compound treatment, cell images, and the true phenotype. The SCM is integrated into the training process to achieve confounder-awareness in our foundation model. By incorporating known confounders into the learning process, the SCM enables the generation of synthetic CP images that account for these biases. Debiasing of subprofile analysis is then achieved through a method adapted from g-estimation (right panel of Figure 2c) [29].

Specifically, in Figures 2b and 2c, we define images O, treatments T (i.e., compounds), and confounders C, as experimental observations. The underlying phenotype P is an unobserved variable which can be estimated from the acquired images O. The images O are acquired via cell microscopy across multiple sources, introducing unavoidable experimental biases, collectively denoted as confounders C. These confounders encompass a range of technical variations inherent to the experimental process, including source, batch, plate and well-position effects. While treatments T should be the primary drivers of phenotypic changes P (indicated with the arrow from T to P), the observed images O provide only a partial and noisy representation of the true phenotypic outcome (indicated with the arrow from P to O) [5, 15]. This necessitates the acquisition of images from hundreds of cells with multiple replicates to enhance the fidelity of phenotypic measurements and mitigate the impact of noise [15]. We hypothesize that our confounder-aware (SCM-conditioned) foundation model, by learning generalizable representations across experimental replicates, can further mitigate this noise and improve the accuracy of phenotypic characterization.

Moreover, experimental and technical variations due to source, batch, plate and well-position (e.g, microscope settings, imaging artifacts, assay preparation) represented by C, exert a multifaceted influence on this causal pathway and directly impact the acquired images (indicated with the arrow from C to O). Concurrently, inherent cell-to-cell variability and variations in cell density or other source-specific experimental conditions influence cellular phenotypes, impacting both cell growth and response (indicated with the arrow from C to P). The potential for systematic bias in treatment allocation is compounded by considerable variations across sources, which may involve different compounds, and by limitations in plate map design (represented by the arrow from C to T). As plate maps are often not fully randomized and may group treatments in specific well positions, these factors, which may vary considerably from source to source, can collectively introduce unintended biases in the experimental process [5, 6, 15]. Hence, our foundation model is further strengthened by the inclusion of a SCM. This SCM is specifically designed to adjust for the known confounders, allowing to debias the downstream analysis at the inference phase, using a method adapted from g-estimation (see the next subsection) [29].

Given the indirect nature of phenotypic observations and the presence of confounding factors, robust cell image analysis is crucial to accurately recover and quantify the effects of T on P. A previous work has implicitly utilized a similar causal interpretation, albeit without explicit formalization [15]. The authors utilized an approach involving implicit causal graph description and batch correction to address confounding factors in treatment classification [15]. Our aim is to disentangle the true biological signal from the unwanted technical variation embedded in the acquired images, thereby enabling accurate inference of compound effects. We introduce a foundation model designed to learn generalizable representations across experimental replicates, able to capture and mitigate various sources of inherent noise and variability. This enhances the accuracy of phenotypic characterization and contributes to the model’s robustness and generalizability. We further augment this foundation model with a SCM to explicitly account for known confounders: i.e., source, batch, plate, and well-position effects. By incorporating the SCM, we aim to generate synthetic CP images that account for known confounders (source, batch and well-position). This leads to more reliable and accurate identification of biological mechanisms and insights.

### Debiasing via a g-estimation-based method and latent diffusion modeling

After modeling the effects of confounders in image generation through confounder-aware (SCM-conditioned) foundation model training (Figure 1a), we are interested in isolating the average effect of a compound T = t on a cell phenotype P during foundation model inference (Figure 1b). The t denotes individual compound treatments, each embedded in the model as a SMILES (1D chemical structure) string. This is equivalent to measuring the expectation of the interventional distribution, E[p(P | do(T = t))]. In practice, we estimate the effect of t on images O via p(O | do(T = t)). We then estimate P from synthetic O, using CellProfiler for cell feature extraction (numerous features relevant to cellular size, shape, intensity and texture across the CP stains) [7] and Subprofile analysis (see relevant subsections later in Methods) [39].

Notably, we estimate the interventional distribution p(O | do(T = t)) via confounder adjustment, also known as the g-estimation [24, 29]. Considering the SCM (DAG) graph associated with the real distribution p(O,T,P,C) learned during confounder-aware (SCM-conditioned) foundation model training, we achieve confounder adjustment at inference by controlling for C, using the backdoor criterion [29]. Specifically, we adjust for confounders C to estimate p(O| do(T=t)), by blocking any backdoor paths from T to O via adjusting for C. Due to the high dimensionality of the images O, traditional confounder adjustment methods that integrate over all C combinations are computationally prohibitive. Therefore, to address this, we utilize Monte Carlo sampling combined with generative (latent diffusion) modeling to approximate the g-estimation and debias our downstream analyses (MoA and target identification). The confounder adjustment at inference can be expressed using the g-estimation formula, based on the backdoor criterion [24, 29]:

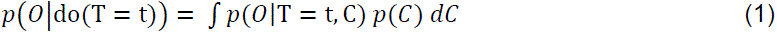

Since we have explicit information about C, we can approximate the integral in Equation (1) by averaging over a range of confounding factors C, using Monte Carlo sampling:

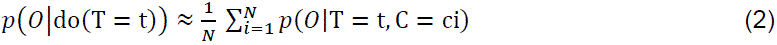

To model the complex distribution p(O | T, C), we employ a conditional generative model in the form of a LDM [40], conditioned on T and C. Diffusion models are a class of deep generative models that synthesize data by reversing a gradual noising process. Trained by systematically corrupting data with Gaussian noise and then learning to denoise, these models can generate high-quality samples from complex distributions, exhibiting remarkable capabilities in image generation [40–42].

By combining LDM with Monte Carlo sampling, we can effectively estimate the interventional distribution p(O | do(T = t)), in the presence of high-dimensional, unstructured data (CP images). This approach allows us to isolate the causal effect of the treatment T on the image O, while accounting for confounding variables C, and adhering to the causal assumptions outlined in our causal diagram. This methodology aligns with the concept of constructing a “fair” synthetic interventional distribution by employing a surrogate do-operation on the conditional distribution [43].

In practice, we synthesize 5-channel CP images across a range of confounding factors ci for a given compound t, i.e., sets of different combinations of source, batch and well-position confounders. ci are sampled from a uniform distribution over the confounder values, enforcing independence between C and T. However, the averaging from Equation 2 does not happen at the image O level, but after feature extraction. We compute the features P (as detailed in “Cell profile extraction” subsection) for each 5-channel CP image. Then, we average the resulting collections of phenotypic cell features, P, where each collection corresponds to a set of 5-channel CP images, generated under a specific set of confounding factors. The algorithmic process is outlined in the Algorithm pseudocode below:

**Algorithm:**
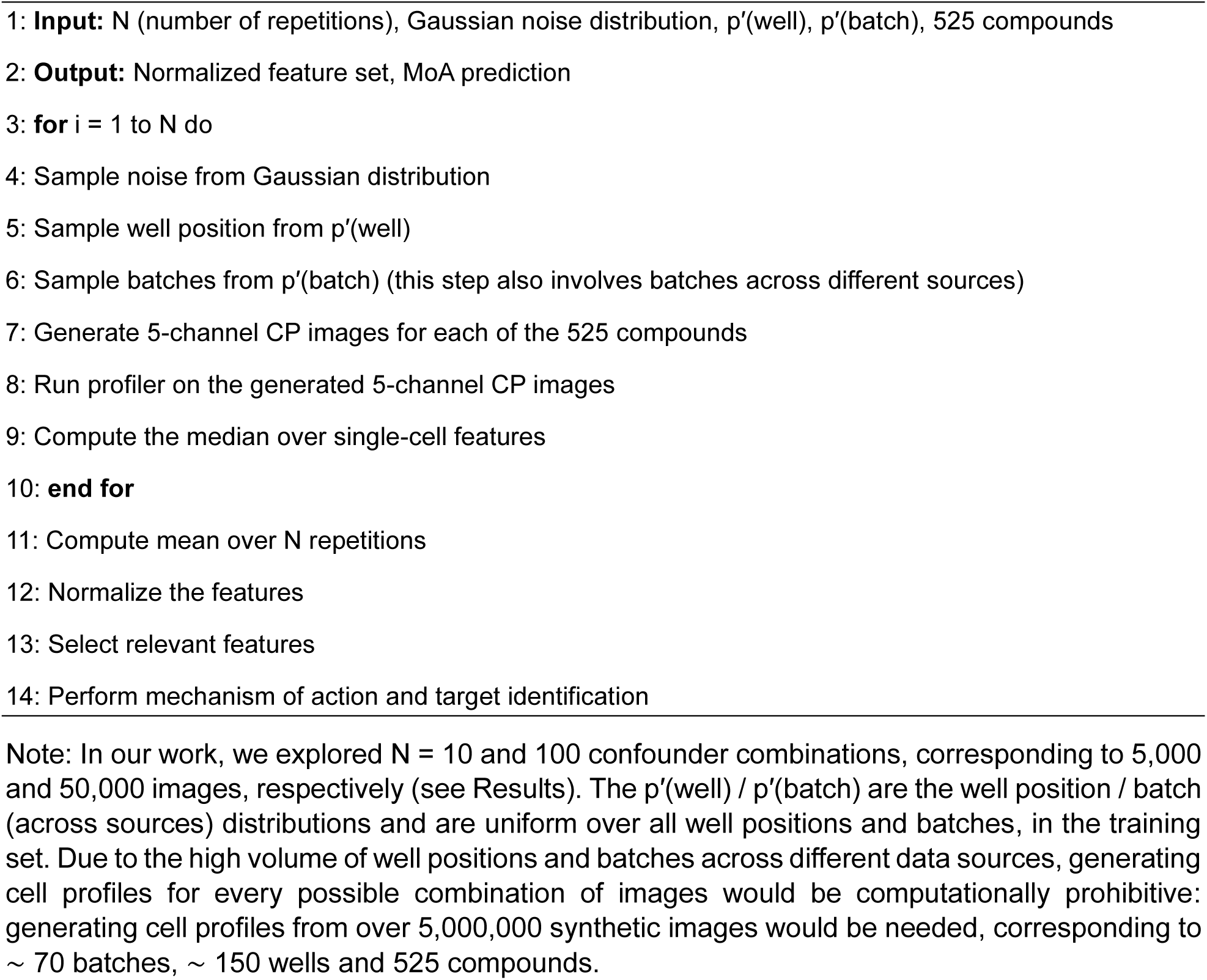
Generation of Synthetic Images.

### Conditional latent diffusion modeling

#### End-to-end confounder-aware LDM framework

The entire method is presented in Figure 1. We developed and trained a confounder-aware foundation model using a LDM conditioned on a SCM, to learn the distribution of cell images [40–42]. LDMs offer a powerful framework for generative modeling, particularly for high-dimensional data such as images. These models operate by first learning a compressed representation of the data in a lower-dimensional latent space [40]. This compression is achieved through a variational autoencoder, which consists of an encoder that maps the input data to the latent space and a decoder that reconstructs the original data from the latent representation [40]. Subsequently, a diffusion model is trained to generate data within this compressed latent space, enabling efficient sampling and manipulation. Our approach generates synthetic CP images based on compound SMILES inputs, while controlling for known confounders via the SCM conditions: source (laboratory), batch and well-position. During training, the confounder-aware model learns to disentangle the effects of known confounders on CP images, effectively mapping confounder combinations (of sources, batches and well-positions) to distinct image semantics. Here, “image semantics” encompasses both cell morphology and broader imaging properties, including intensity, texture, spatial arrangement, and fluorescence, which can all be influenced by confounders. To mitigate the influence of confounding factors during inference, we employed a g-estimation-based approach based on Monte Carlo sampling (as described above), to generate CP images across a diverse range of source, batch, and well-position combinations [29]. By then averaging the extracted cell profiles from these images, we effectively controlled for the impact of these confounders, thereby debiasing the downstream tasks of MoA and target identification.

Our confounder-aware (SCM-conditioned) foundation model was designed to generalize beyond the JUMP-CP training data and accurately estimate the effects of novel compounds. To achieve this, we employed a pretrained MolT5 encoder [28], as described in the next subsection. This encoder, trained on a vast dataset of over 200 million molecules in SMILES format (Figure 1), generated robust chemical representations, independently of the associated images. The integration of MolT5-derived embeddings as conditioning factors, allowed our LDM to effectively learn the mapping between molecular information and CP image semantics, enabling accurate image generation for both seen and unseen compounds.

#### Architecture and training of the LDM

In our work, we utilized the pretrained variational autoencoder from the Stable Diffusion XL method [44], to obtain latent representations of our cell images. This autoencoder was originally trained on natural images with 3 (RGB) channels, exhibiting a compression rate of 8x for each spatial dimension. To accommodate the 5-channel nature of our cell images, which exceeds the 3-channel input capacity of the Stable Diffusion XL autoencoder [44], we employ a channel-wise encoding strategy. Specifically, we divide the image channels into two groups (each with 3-channel inputs, one of the channels repeating between groups), encode each group independently using the pretrained autoencoder, and subsequently concatenate the resulting latent representations. This yields a final latent representation with 8 channels, which can effectively capture the information content of our cell images.

To facilitate the manipulation of our latent representations, we constructed a diffusion model with a U-Net architecture [45] based on the original LDM implementation [41]. However, in contrast to utilizing a pretrained model, we opted to train our diffusion model from scratch. The U-Net architecture comprises 3 levels, each with 2 residual blocks [46] for efficient feature learning. The number of channels in the convolutional layers within the U-Net increases progressively across the 3 levels, with 256, 512, and 768 channels, respectively. Furthermore, to capture long-range dependencies within the latent representations, attention blocks were integrated into the 2 deepest levels of the U-Net [41].

We used the MONAI generative models package for developing, training and testing our SCM-conditioned (confounder-aware) and SCM-free foundation LDMs [47]. Our CP images underwent preprocessing where, for each channel (stain), pixel values were normalized to a range of [-1, 1]. Subsequently, we trained our models on centrally cropped regions of these images, each with a dimension of 384 × 384 pixels. We trained our diffusion model for 30 epochs with a batch size of 192 images, utilizing the AdamW optimizer [48]. For the diffusion noise schedule, we employed a scaled linear profile with 1000 diffusion steps. The noise levels, defined by beta values, ranged from 0.0015 to 0.0205. The model was trained with angular parameterization (also known as v-prediction) which focuses on predicting a mixed representation of the image and the noise at each time-step, within the LDM framework [49]. Employing the angular representation for training diffusion models has been shown to yield a more stable objective function [49], compared to the original approaches [40]. The training of the LDM was performed using 4 V100 GPUs with 40Gb of memory each.

### Encoding compounds via a Transformer encoder

A critical aspect of our methodology involves establishing distinct embedding profiles for each chemical compound. These profiles must robustly and consistently capture the similarities and differences in chemical structures across the 107,289 compounds used to train the LDM. This is crucial because each compound’s embedding serves as a separate conditioning factor, enabling the disentanglement of subtle and evident variations among compounds during image generation.

Of note, a key objective of our SCM-conditioned foundation model is to accurately estimate the effects of novel compounds, i.e., those entirely unseen during training. This generalization capability necessitates that the diffusion model effectively learns and encodes the similarities and differences within the molecular representations of the training compounds. To achieve this, we leverage the widely adopted SMILES representation for encoding molecular structures. We then encode these SMILES representations using a pretrained self-supervised learning framework, MolT5 (Molecular T5), which has demonstrated strong performance in capturing molecular features [28]. MolT5 has demonstrated a remarkable ability to generate captions (chemical structure descriptions) from molecular information (SMILES) and to create a molecule (SMILES) that matches a given natural language description. This proficiency stems from MolT5’s pretraining on the vast amount of unlabeled natural language text and molecule strings from the ZINC-15 dataset, which contains approximately 200 million molecules with SMILES information [50].

We employed this pretrained MolT5 model to derive embeddings for each compound, which were then used as conditioning factors during both the LDM training and inference stages.

### Cell profile extraction

Cell profiles, organized by batch and source, were obtained from the JUMP-CP data gallery [2], for all real CP data used to train the LDM. These profiles were derived using CellProfiler, an open-source image analysis software for extracting quantitative features from microscopy images. We employed the same software (version 4) to calculate cell profiles for the confounder-aware (SCM-conditioned) and non-confounder-aware (SCM-free) model-derived synthetic CP images, ensuring consistency in feature extraction across all datasets [7]. This software facilitates the construction of automated pipelines for high-throughput image analysis, enabling the measurement of various cellular properties such as size, shape, intensity, and texture [15, 39]. The CellProfiler image processing workflow comprises 3 sub-tasks: (1) illumination correction, (2) quality control, and (3) feature extraction.

Non-homogeneous illumination across the image field, a common artifact in high-throughput microscopy, introduces systematic bias and can lead to measurement errors. To mitigate these effects, illumination correction is first performed successively on each channel (DNA, ER, AGP, and Mito). This module employs a median filter to approximate the illumination distribution across the image [51].

The second step encompasses quality control and the application of illumination correction. Initially, image quality is assessed across all channels using metrics such as blur measurements, saturation, and intensity [52]. Images failing the predefined quality criteria are flagged and excluded from further analysis. Finally, each image channel is illumination corrected [51].

The primary feature extraction pipeline involves segmenting the images to identify objects of interest: nuclei, cells, and cytoplasm. Initially, nuclei are identified based on the DNA image using advanced settings optimized for precise boundary detection and separation of closely spaced nuclei. This identification employs a global Otsu-based thresholding method for initial segmentation. Subsequently, cell segmentation is performed by expanding regions around the identified nuclei, using the Watershed algorithm [53].

Following cell segmentation, the CellProfiler extracts single-cell features, which are designed to be human readable and grouped by cell region (nucleus, cytoplasm or cell) [7, 51]. Overall, 5,797 features are extracted from the 3 cell regions that aim to characterize the effect of the screened compound on the cell line at different levels. Individual single-cell-derived features are then combined across all cells at the image level, using the median value [51]. Next, features from individual images (fields of view) are averaged to create a well-level profile. Finally, treatment-level profiles are obtained by averaging all available replicates of wells (across plates).

### Batch correction of real JUMP-CP data

To mitigate batch effects in the real JUMP-CP data and perform a fair comparison against our SCM-conditioned foundation model, we employed the Harmony algorithm for batch correction [54]. Harmony is a batch effect correction technique developed primarily for correcting batch effects in scRNAseq data and has recently been adopted for image-based cell profiles [21, 54].

The method employs an iterative approach to learn a cell-profile-specific linear transformation. It alternates between two steps: (1) maximizing the diversity of fuzzy clusters with respect to batch variations, and (2) correcting batch effects by applying a mixture model. A limitation of the Harmony algorithm is its requirement to recompute the linear transformations across the entire dataset each time a new profile is introduced. However, a recent comprehensive evaluation of batch correction methods for CP data, demonstrated that Harmony exhibits leading batch effect correction performance on profiles derived from real images, such as those from the JUMP-CP dataset [21]. Harmony is a widely used and effective method for batch effect correction, commonly employed to minimize the effects of known biases in real CP data [21].

### MoA and target identification via subprofiling

#### Calculating “reference subprofiles” and Biosimilarity

Generating CP profiles for reference compounds, which are characterized for their MoA through bioactivity assays and other complementary approaches, is an important prerequisite for analyzing the bioactivity of novel or uncharacterized chemical compounds [5]. Ideally, reference compounds sharing a common MoA or target should exhibit similar CP profiles, enabling MoA/ target prediction for novel compounds based on profile Biosimilarity. However, this approach can be confounded by incomplete annotation or polypharmacology exhibited by reference compounds, necessitating a more nuanced analysis [39]. Subprofiling, a recently developed technique for fast and precise prediction of MoA characterization, was applied in our study [39].

We used subprofile analysis for both MoA, and for the first-time, for target identification. This approach involves the definition of “reference subprofiles”, which are subsets of features common to reference compounds within a single MoA-or target-specific cluster. It is anticipated that comparing the similarity of a new compound’s cell profile to a reference MoA-or target-specific cluster subprofile, will substantially improve and accelerate the MoA/ target identification of novel or uncharacterized compounds [39]. Beginning with the full profiles of a set of MoA-or target-specific reference compounds, dominant features are first extracted. A representative consensus subprofile is then defined, to encapsulate the properties of the set. The consensus subprofile is the median of the cluster, which we name here as “reference subprofile” [39].

Specifically, for a group of reference compounds sharing a common MoA/ target, each measured feature within the full profile is evaluated. A feature is retained in the final subprofile only if it exhibits a consistent directional response (either positive or negative) across a significant majority (greater than 85%) of the profiles within the cluster. This process culminates in the selection of subprofile features, with their final values determined as the median values across all compound profiles within the group, i.e., the “reference subprofile”. The “reference subprofile” can then be employed to calculate the Biosimilarity of novel or uncharacterized compounds. In subprofiling, the Biosimilarity between two profiles, i.e., a novel or uncharacterized compound and a “reference subprofile”, (u, v) is defined as:

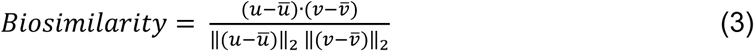

where the 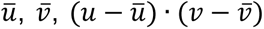 and 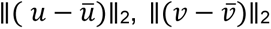 define the mean of the considered profiles (vectors) u and v, the dot product and the Euclidean norms of the considered profiles, respectively.

#### Implementation

In our work, Biosimilarity analysis was employed to quantitatively assess the confounder-aware– and non-confounder-aware foundation model performance, against real and real-batch-corrected JUMP-CP data, in the downstream application of MoA and target identification. To perform this evaluation, from the 107,289 compounds used to train our model, we selected those with annotations for both MoAs and targets in the Broad Institute Drug Repurposing Hub. Supplementary Table 2 presents the number of compounds used for biological effect estimation (MoA and target prediction) across each fold.

To ensure balanced representation in our Biosimilarity analysis, we further filtered this selection to include only MoAs and targets associated with at least 10 compounds (Supplementary Figure 3). To minimize polypharmacology effects in our MoA reference subprofiles, we further refined our selection to include only compounds with a single associated MoA. This resulted in a final dataset of 525 and 465 compounds for MoA and target identification, respectively. This dataset encompassed 26 distinct MoAs and 446 distinct targets (there could be multiple targets per compound).

For each MoA and target, we randomly sampled 5 compounds to generate “reference subprofiles”. These subprofiles serve as baseline activity patterns for comparison. This process yielded 130 compounds with MoA annotations, covering 26 distinct MoAs, which were used to establish MoA-based reference subprofiles. For target-based reference subprofiles, we varied the number of compounds from 88-98, depending on the classification fold (see “Evaluation metrics”). The remaining 395 compounds and 367-377 compounds, out of the 525 and 465 respectively, were used to perform Biosimilarity analysis for the confounder-aware– and non-confounder-aware foundation model-derived synthetic images. To evaluate the ability of Biosimilarity scores to identify the correct MoA, we used 273-281 compounds seen during training and 111-122 unseen compounds (out of a total of 395). Similarly, for target identification, we used 228-238 compounds seen during training and 139 unseen compounds (out of 367-377 total compounds). Subprofile extraction was performed using a Python implementation adapted from the authors’ publicly available repository (https://github.com/mpimp-comas/2022_pahl_ziegler_subprofiles) [39]. This implementation served as a reference for extracting subprofiles from the SCM-conditioned and SCM-free foundation model-derived synthetic images, as well as the real and real batch-corrected images from the JUMP-CP dataset.

### Evaluation metrics

To evaluate the accuracy of the estimated Biosimilarities in identifying known MoA and target annotations, we devised a nearest neighbor classifier. This evaluation leveraged ground truth MoA and target annotations for compounds from the Drug Repurposing Hub. To assess the performance of our nearest neighbor classifier in identifying MoA and target ground truths, we employed mean Average Precision (mAP) [30] and area under the receiver operating characteristic curve (ROC-AUC) [31], as evaluation metrics. These metrics are well-suited for evaluating biological effect prediction, particularly when prediction thresholds for true positive rate (TPR) and false positive rate (FPR) may typically differ between synthetic and real data. The formulation of each metric is detailed below.

mAP evaluates the model’s ability to correctly rank relevant labels (here: known MoA or target annotations). It summarizes the precision-recall curve by calculating the weighted mean of precisions at each recall level, with the weight being the increase in recall from the previous threshold. For each sample, the average precision (AP) is defined as:

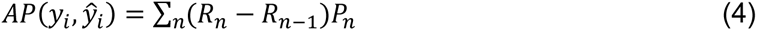

where 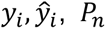 and R_*n*_ are the true value, predicted value, precision and recall at the n-th threshold, respectively. The mAP aggregates the AP values across all samples as:

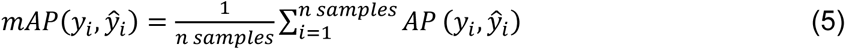

A higher mAP score indicates better performance, with a perfect score of 1 achieved when all relevant labels are ranked before any irrelevant ones across all samples. This metric is particularly suitable for evaluating multi-label classification tasks (here: Biosimilarity scores against multiple MoA and target “reference subprofiles”), as it accounts for the order of relevant labels and balances precision at varying recall levels.

The area under the receiver operating characteristic curve ROC-AUC evaluates the model’s classification accuracy by measuring the trade-off between the true positive rate (TPR) and the false positive rate (FPR), across all possible classification thresholds. It provides a single scalar value that summarizes the model’s performance over the entire range of operating conditions, effectively capturing its discriminative capability.

For each label in a multi-label classification task, the ROC-AUC is calculated by plotting the TPR against the FPR at various threshold settings. The TPR and FPR are defined as:

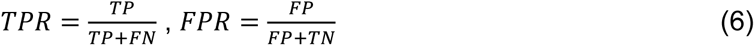

where TP, FP, TN, and FN represent the counts of true positives, false positives, true negatives, and false negatives, respectively.

The ROC-AUC for each label is calculated as the area under its corresponding ROC curve, typically using the trapezoidal rule for numerical integration. In a multi-label setting, the overall ROC-AUC is obtained by averaging the ROC-AUC values across all labels:

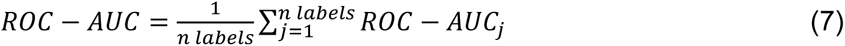

where R*OC* − *AUC*_*j*_ denotes the ROC-AUC for the j-th label. A ROC-AUC value of 1 indicates perfect classification, with the model correctly distinguishing between all positive and negative instances. An ROC-AUC of 0.5 suggests no discriminative ability, equivalent to random guessing. Therefore, higher ROC-AUC scores reflect better model performance. This metric is particularly suitable for evaluating models on imbalanced datasets, as it is insensitive to the label distribution and focuses on the ranking quality of the predictions rather than their absolute values. By considering all classification thresholds, ROC-AUC provides a comprehensive assessment of the model’s ability to prioritize relevant labels over irrelevant ones.

## Data Availability

The synthetic image and profile datasets, along with qualitative results and numeric values for metrics is available at http://dmjump-sinkove-pfizer-results.s3-website-us-east-1.amazonaws.com. Ground truth for the experiments of MoA and target prediction were collected from the drug repurposing hub at https://repo-hub.broadinstitute.org/repurposing-app.

## Acknowledgements

This study was sponsored by Pfizer. The sponsor played no role in study design, data collection, analysis and interpretation of data, or the writing of this manuscript. We are grateful to Dr. Regis Doyonnas, Dr. David Logan and Dr. Vladimir Ivanov from Pfizer, for their valuable discussions and feedback during the initial stages of our method development.

## Author contributions statement

GP: author of the manuscript. Conceptualized the objectives and study design, conceived and conducted the experiments, devised the methods, performed analysis, designed and developed the analysis software and interpreted the findings of the manuscript. PS: author of the manuscript. Conceptualized and devised the methods, conceived and conducted the experiments, performed analysis, designed and developed the analysis software. AC: devised the methods, conducted the experiments, performed analysis, developed the analysis software. GY: contributed to the evaluation of the methods, interpreted the findings, edited the manuscript. WHLP: conducted the experiments, performed analysis, developed the analysis software. All authors reviewed the manuscript.

## References

1. Chandrasekaran, S. N. et al. Three million images and morphological profiles of cells treated with matched chemical and genetic perturbations. Nat. Methods 1–8 (2024).

2. Weisbart, E. et al. Cell painting gallery: an open resource for image-based profiling. Nat. Methods 1775–1777 (2024).

3. Bray, M.-A. et al. Cell painting, a high-content image-based assay for morphological profiling using multiplexed fluorescent dyes. Nat. protocols 11, 1757–1774 (2016).

4. Ziegler, S., Sievers, S. & Waldmann, H. Morphological profiling of small molecules. Cell chemical biology 28, 300–319 (2021).

5. Chandrasekaran, S. N., Ceulemans, H., Boyd, J. D. & Carpenter, A. E. Image-based profiling for drug discovery: due for a machine-learning upgrade? Nat. Rev. Drug Discov. 20, 145–159 (2021).

6. Chandrasekaran, S. N. et al. Jump cell painting dataset: morphological impact of 136,000 chemical and genetic perturbations. BioRxiv 2023–03 (2023).

7. Stirling, D. R. et al. Cellprofiler 4: improvements in speed, utility and usability. BMC bioinformatics 22, 1–11 (2021).

8. Kraus, O. Z., Ba, J. L. & Frey, B. J. Classifying and segmenting microscopy images with deep multiple instance learning. Bioinformatics 32, i52–i59 (2016).

9. Pawlowski, N., Caicedo, J. C., Singh, S., Carpenter, A. E. & Storkey, A. Automating morphological profiling with generic deep convolutional networks. BioRxiv 085118 (2016).

10. Janssens, R., Zhang, X., Kauffmann, A., de Weck, A. & Durand, E. Y. Fully unsupervised deep mode of action learning for phenotyping high-content cellular images. Bioinformatics 37, 4548– 4555 (2021).

11. Perakis, A. et al. Contrastive learning of single-cell phenotypic representations for treatment classification. In Machine Learning in Medical Imaging: 12th International Workshop, MLMI 2021, Held in Conjunction with MICCAI 2021, Strasbourg, France, September 27, 2021, Proceedings 12, 565–575 (Springer, 2021).

12. Cross-Zamirski, J. O. et al. Self-supervised learning of phenotypic representations from cell images with weak labels. In NeurIPS 2022 Workshop on Learning Meaningful Representations of Life (2022).

13. Wong, D. R. et al. Deep representation learning determines drug mechanism of action from cell painting images. Digit. Discov. 2, 1354–1367 (2023).

14. Moshkov, N. et al. Predicting compound activity from phenotypic profiles and chemical structures. Nat. communications 14, 1967 (2023).

15. Moshkov, N. et al. Learning representations for image-based profiling of perturbations. Nat. communications 15, 1594 (2024).

16. Lamiable, A. et al. Revealing invisible cell phenotypes with conditional generative modeling. Nat. communications.14, 6386 (2023).

17. Christiansen, E. M. et al. In silico labeling: predicting fluorescent labels in unlabeled images. Cell 173, 792–803 (2018).

18. Lee, G., Oh, J.-W., Her, N.-G. & Jeong, W.-K. Deephcs++: Bright-field to fluorescence microscopy image conversion using multi-task learning with adversarial losses for label-free high-content screening. Med. Image analysis 70, 101995 (2021).

19. Cross-Zamirski, J. O. et al. Label-free prediction of cell painting from brightfield images. Sci. reports 12, 10001 (2022).

20. Xing, X. et al. Can generative ai replace immunofluorescent staining processes? a comparison study of synthetically generated cellpainting images from brightfield. Comput. Biol. Medicine 182, 109102 (2024).

21. Arevalo, J. et al. Evaluating batch correction methods for image-based cell profiling. Nat. Commun. 15, 6516 (2024).

22. Pawlowski, N., Coelho de Castro, D. & Glocker, B. Deep structural causal models for tractable counterfactual inference. Adv. neural information processing systems 33, 857–869 (2020).

23. Sanchez, P. & Tsaftaris, S. A. Diffusion causal models for counterfactual estimation. In First Conference on Causal Learning and Reasoning (2022).

24. Pearl, J. Causality (Cambridge university press, 2009).

25. Koller, D. & Friedman, N. Probabilistic graphical models: principles and techniques (MIT press, 2009).

26. Schölkopf, B. et al. Toward causal representation learning. Proc. IEEE 109, 612–634 (2021).

27. Melistas, T., et al. Benchmarking counterfactual image generation. In The Thirty-eight Conference on Neural Information Processing Systems Datasets and Benchmarks Track (2024).

28. Edwards, C. et al. Translation between molecules and natural language. In 2022 Conference on Empirical Methods in Natural Language Processing, EMNLP 2022 (2022).

29. Robins, J. A new approach to causal inference in mortality studies with a sustained exposure period—application to control of the healthy worker survivor effect. Math. modelling 7, 1393–1512 (1986).

30. Manning, C. D., Raghavan, P. & Schütze, H. Boolean retrieval. Introd. to information retrieval 1–18 (2008).

31. Fawcett, T. An introduction to roc analysis. Pattern recognition letters 27, 861–874 (2006).

32. Hochberg, Y. Multiple comparison procedures (1987).

33. Vincent, F. et al. Hit triage and validation in phenotypic screening: considerations and strategies. Cell Chem. Biol. 27, 1332–1346 (2020).

34. Sanchez-Fernandez, A., Rumetshofer, E., Hochreiter, S. & Klambauer, G. Cloome: contrastive learning unlocks bioimaging databases for queries with chemical structures. Nat. communications. 14, 7339 (2023).

35. Xing, X. et al. Artificial immunofluorescence in a flash: Rapid synthetic imaging from brightfield through residual diffusion. Neurocomputing 612, 128715 (2025).

36. Fay, M. M. et al. Rxrx3: Phenomics map of biology. Biorxiv 2023–02 (2023).

37. Corsello, S. M. et al. The drug repurposing hub: a next-generation drug library and information resource. Nat. medicine 23, 405–408 (2017).

38. Niepel, M. et al. A multi-center study on the reproducibility of drug-response assays in mammalian cell lines. Cell systems 9, 35–48 (2019).

39. Pahl, A. et al. Morphological subprofile analysis for bioactivity annotation of small molecules. Cell Chem. Biol. 30, 839–853 (2023).

40. Ho, J., Jain, A. & Abbeel, P. Denoising diffusion probabilistic models. Adv. neural information processing systems 33, 6840–6851 (2020).

41. Rombach, R., Blattmann, A., Lorenz, D., Esser, P. & Ommer, B. High-resolution image synthesis with latent diffusion models. In Proceedings of the IEEE/CVF conference on computer vision and pattern recognition, 10684–10695 (2022).

42. Dhariwal, P. & Nichol, A. Diffusion models beat gans on image synthesis. Adv. neural information processing systems 34, 8780–8794 (2021).

43. Van Breugel, B., Kyono, T., Berrevoets, J. & Van der Schaar, M. Decaf: Generating fair synthetic data using causally-aware generative networks. Adv. Neural Inf. Process. Syst. 34, 22221–22233 (2021).

44. Podell, D., et al. Sdxl: Improving latent diffusion models for high-resolution image synthesis. In The Twelfth International Conference on Learning Representations (2024).

45. Ronneberger, O., Fischer, P. & Brox, T. U-net: Convolutional networks for biomedical image segmentation. In Medical image computing and computer-assisted intervention–MICCAI 2015: 18th international conference, Munich, Germany, October 5-9, 2015, proceedings, part III 18, 234–241 (Springer, 2015).

46. He, K., Zhang, X., Ren, S. & Sun, J. Deep residual learning for image recognition. In Proceedings of the IEEE conference on computer vision and pattern recognition, 770–778 (2016).

47. Pinaya, W. H. et al. Generative ai for medical imaging: extending the monai framework. arXiv preprint arXiv:2307.15208 (2023).

48. Loshchilov, I. & Hutter, F. Decoupled weight decay regularization. In International Conference on Learning Representations (2019).

49. Salimans, T. & Ho, J. Progressive distillation for fast sampling of diffusion models. In International Conference on Learning Representations (2022).

50. Sterling, T. & Irwin, J. J. Zinc 15–ligand discovery for everyone. J. chemical information modeling 55, 2324–2337 (2015).

51. Singh, S., Bray, M.-A., Jones, T. & Carpenter, A. Pipeline for illumination correction of images for high-throughput microscopy. J. microscopy 256, 231–236 (2014).

52. Bray, M.-A., Fraser, A. N., Hasaka, T. P. & Carpenter, A. E. Workflow and metrics for image quality control in large-scale high-content screens. J. biomolecular screening 17, 266–274 (2012).

53. Vincent, L. & Soille, P. Watersheds in digital spaces: an efficient algorithm based on immersion simulations. IEEE Transactions on Pattern Analysis & Mach. Intell. 13, 583–598 (1991).

54. Korsunsky, I. et al. Fast, sensitive and accurate integration of single-cell data with harmony. Nat. methods 16, 1289–1296 (2019).

