## Supplementary Information for "Confounder-aware foundation modeling for accurate phenotype profiling in cell imaging"

### Supplementary Notes

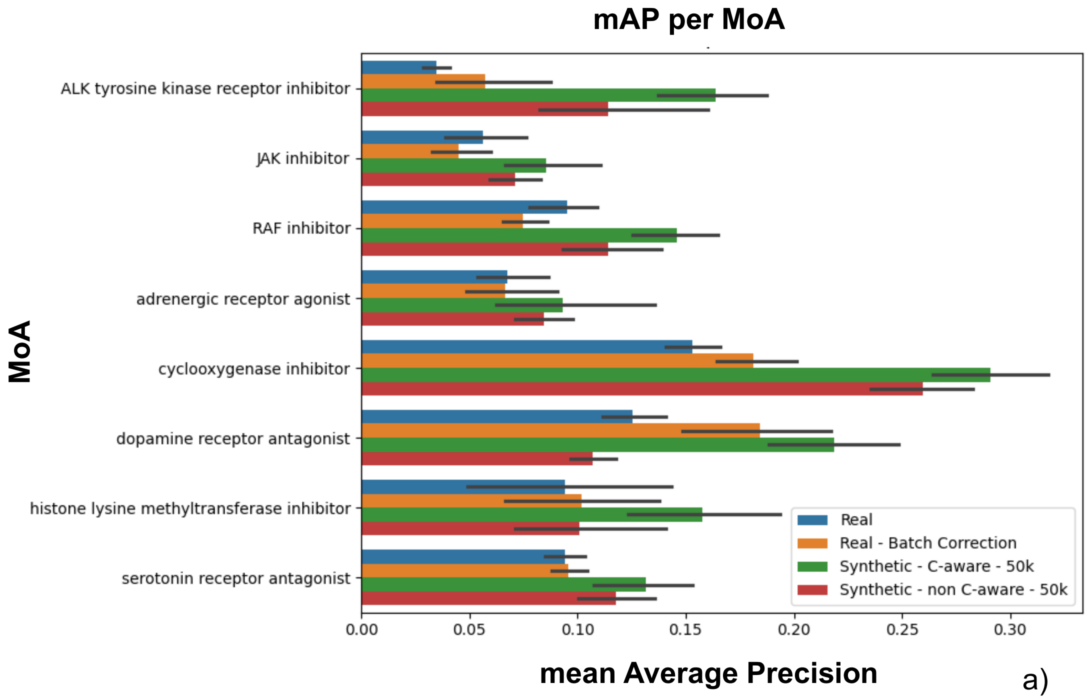


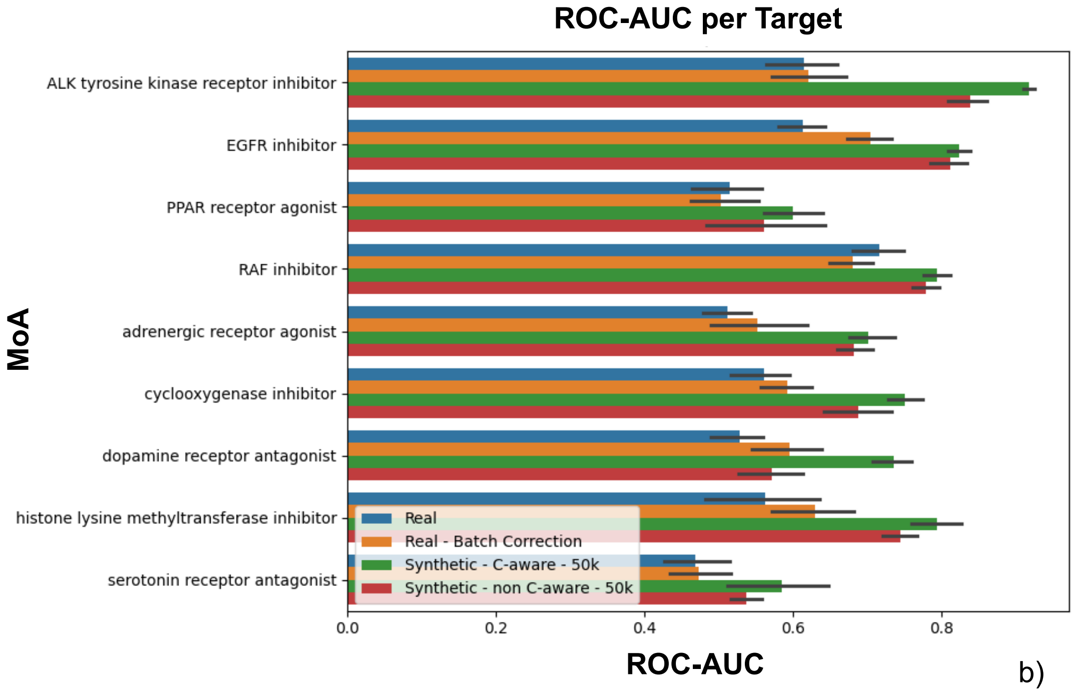


**Supplementary Figure 1)** mAP (mean average precision) (a) and ROC-AUC (b) per MoA, highlighting instances where the confounder-aware model outperformed other approaches.

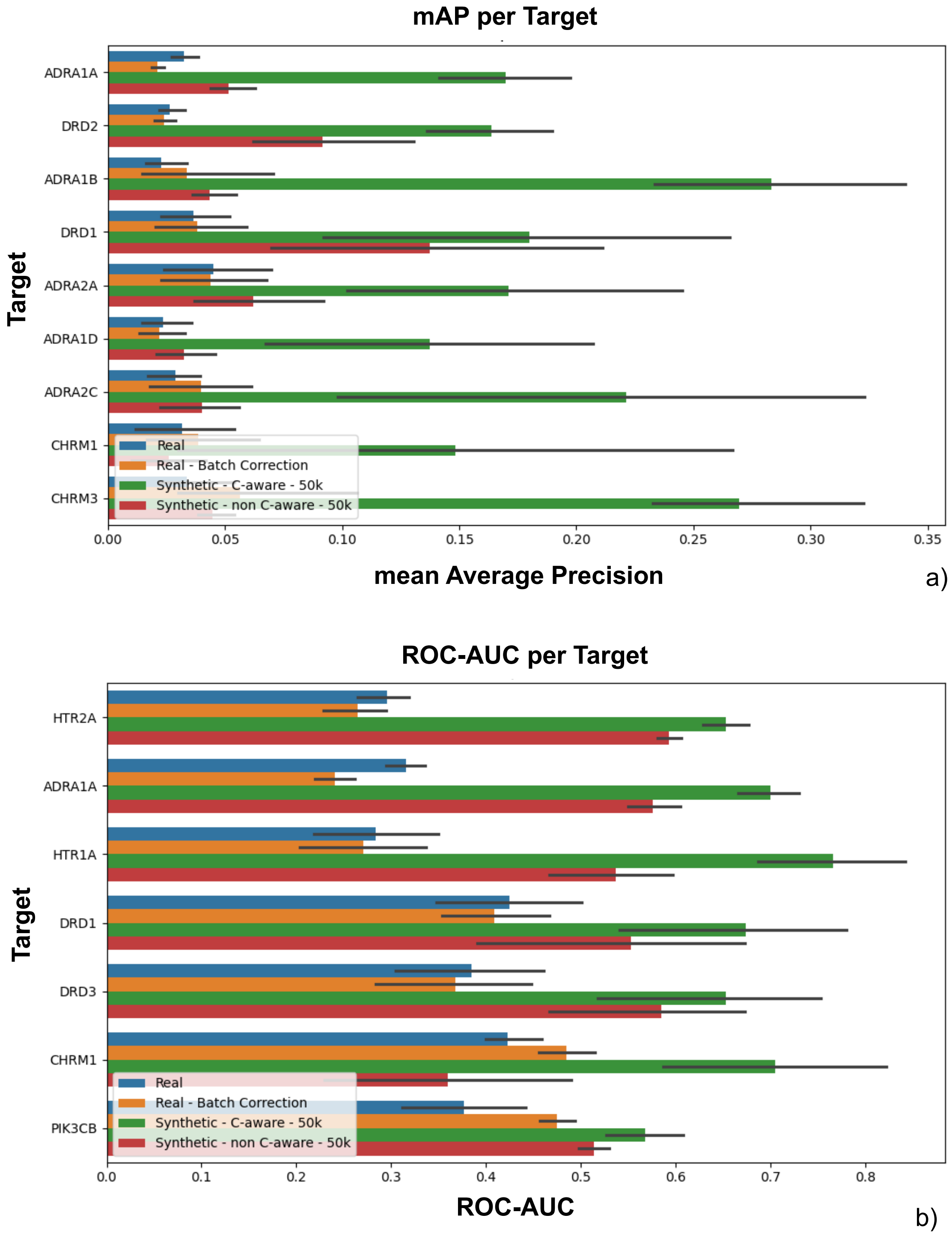


**Supplementary Figure 2)** mAP (mean average precision) and ROC-AUC per compound target, highlighting instances where the confounder-aware model outperformed other approaches.

### Supplementary Methods

**Supplementary Table 1)** Summary of data and experimental setup for evaluating confounder-aware and non-confounder-aware foundation models

| Aspect | Description | Algorithm Settings |
|---|---|---|
| Image source | JUMP-CP Consortium dataset (cpg0016) | |
| Cell line | U2OS | |
| Image type | 5-channel Cell Painting (CP) images | |
| Total images | 13,361,250 CP images (2,672,250 5-channel CP images) | |
| Total (5-channel) image sets per source | Source 1: 292,002<br>Source 2: 443,738<br>Source 3: 809,622<br>Source 9: 583,750<br>Source 11: 543,138 | |
| Compound selection | 107,289 compounds from all 5 sources with partially overlapping compound sets | |
| Data organization | 48 batches, 832 plates (384 to 1536 wells per plate), 4 to 16 fields of view per well | |
| Confounders | Source (laboratory), batch, and well position | |
| Foundation models | Confounder-aware (SCM-conditioned); Non-confounder-aware (SCM-free) | Trained on the principal dataset (cpg0016) |
| Evaluation data | MoA and target annotations from the Broad Institute Drug Repurposing Hub for 525 compounds. Compounds seen and unseen during model training. | |
| Evaluation metrics | Mean Average Precision, Area Under the Receiver Operating Characteristic Curve | |
| Evaluation tasks | MoA prediction, Compound target prediction | Applied to both synthetic images (from seen and unseen compounds: 525 annotated compounds in total) and real images |

**Supplementary Table 2)** Number of compounds used for MoA and target prediction, for each fold and task.

| Task | Fold | Used for evaluation | Used for reference "subprofiles" | Total | Used to evaluate seen compounds | Used to evaluate unseen compounds |
|---|---|---|---|---|---|---|
| MoA | 1 | 395 | 130 | 525 | 281 | 114 |
| MoA | 2 | 395 | 130 | 525 | 280 | 115 |
| MoA | 3 | 395 | 130 | 525 | 278 | 117 |
| MoA | 4 | 395 | 130 | 525 | 278 | 117 |
| MoA | 5 | 395 | 130 | 525 | 279 | 116 |
| MoA | 6 | 395 | 130 | 525 | 273 | 122 |
| MoA | 7 | 395 | 130 | 525 | 277 | 118 |
| MoA | 8 | 395 | 130 | 525 | 284 | 111 |
| MoA | 9 | 395 | 130 | 525 | 278 | 117 |
| MoA | 10 | 395 | 130 | 525 | 281 | 114 |
| Target | 1 | 368 | 97 | 465 | 229 | 139 |
| Target | 2 | 373 | 92 | 465 | 234 | 139 |
| Target | 3 | 370 | 95 | 465 | 231 | 139 |
| Target | 4 | 368 | 97 | 465 | 229 | 139 |
| Target | 5 | 369 | 96 | 465 | 230 | 139 |
| Target | 6 | 367 | 98 | 465 | 228 | 139 |
| Target | 7 | 370 | 95 | 465 | 231 | 139 |
| Target | 8 | 369 | 96 | 465 | 230 | 139 |
| Target | 9 | 367 | 98 | 465 | 228 | 139 |
| Target | 10 | 377 | 88 | 465 | 238 | 139 |

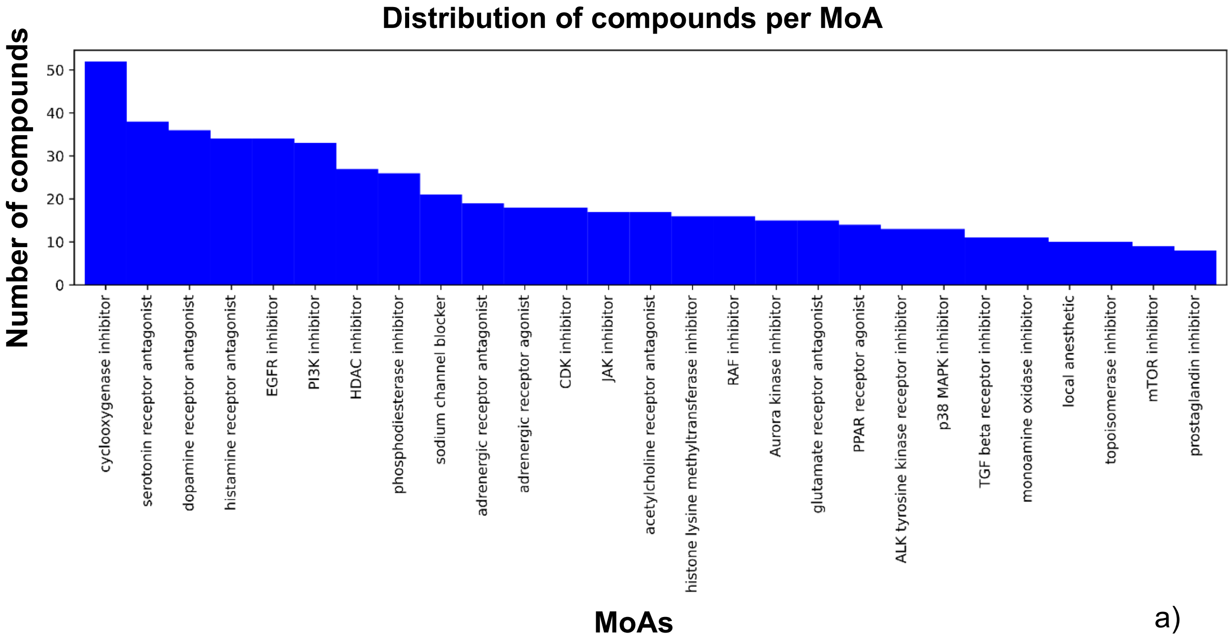


a)

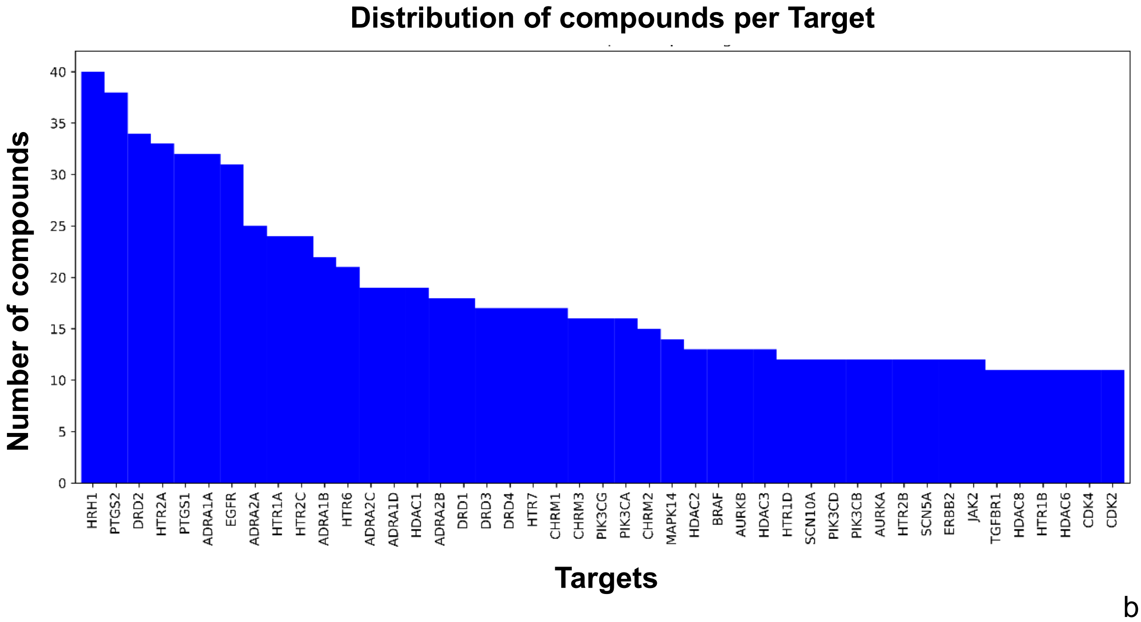


b)

**Supplementary Figure 3)** Distribution of compounds across MoAs and selected compound targets from the Broad Institute Drug Repurposing Hub. MoA: mechanism of action.
